# An adaptive behavioral control motif mediated by cortical axo-axonic inhibition

**DOI:** 10.1101/2023.03.10.531767

**Authors:** Kanghoon Jung, Minhyeok Chang, André Steinecke, Benjamin Berke, Youngjin Choi, Yasuhiro Oisi, David Fitzpatrick, Hiroki Taniguchi, Hyung-Bae Kwon

## Abstract

Neural circuits are reorganized with specificity during learning. Genetically-defined subgroups of inhibitory interneurons are thought to play distinct roles in learning, but heterogeneity within these subgroups has limited our understanding of the scope and nature of their specific contributions to learning. Here we reveal that the chandelier cell (ChC), an interneuron type that specializes in inhibiting the axon-initial segment (AIS) of pyramidal neurons, establishes cortical microcircuits for organizing neural coding through selective axo-axonic synaptic plasticity. We find that organized motor control is mediated by enhanced population coding of direction-tuned premotor neurons, whose tuning is refined through suppression of irrelevant neuronal activity. ChCs are required for learning-dependent refinements via providing selective inhibitory control over pyramidal neurons rather than global suppression. Quantitative analysis on structural plasticity of axo-axonic synapses revealed that ChCs redistributed inhibitory weights to individual pyramidal neurons during learning. These results demonstrate an adaptive logic of the inhibitory circuit motif responsible for organizing distributed neural representations. Thus, ChCs permit efficient cortical computation in a target cell specific manner, which highlights the significance of interneuron diversity.

## Introduction

The reorganization of neural circuits during learning relies on a complex interplay between excitatory and inhibitory signals in the brain, accompanying highly specific synaptic modification. Various synaptic changes contributing to this interplay have been observed at the scale of individual cells and whole circuits. The observed formation and elimination of dendritic spines in cortical pyramidal neurons (PyNs) during learning have suggested that learning is associated with changes in neuronal connectivity ^1–4^. At the circuit level, neuronal activity is synchronized in a subpopulation of neurons, resulting in a high correlation between neuronal activity patterns and learned behaviors ^3, 5, 6^. It has been understood that GABAergic inhibition plays a critical role in shaping learning-dependent circuit changes. Interneurons are categorized by anatomical and electrophysiological features, and their distinct functions during learning have been shown in genetically-defined subtypes such as parvalbumin (PV)-, somatostatin (SOM)- and vasoactive intestinal polypeptide (VIP)-expressing neurons ^7–11^. However, these subtypes still contain a remarkable degree of heterogeneity, such that the precise role of inhibition mediated by single-type interneurons remains an open question.

The ChC (i.e., ‘axo-axonic cell’) is a *bona fide* GABAergic interneuron subclass with respect to its distinct axonal geometry, subcellular synapse connectivity, and fast-spiking electrophysiological properties ^12, 13^. A single ChC exhibits a characteristic axonal geometry with many prominent vertical branches, which contain strings of synaptic boutons exclusively aligned along the AISs of neighboring PyNs. Because the AIS is the site of action potential initiation, ChCs can have decisive control over spike generation in an ensemble of PyNs, thereby regulating synchrony and oscillation of network activity. These functions are thought to be critical for higher-order cognitive processes like working memory ^14–16^. In this study, using a transgenic mouse line specifically engineered to target ChCs, we examined the role of ChCs in sculpting cortical circuits involved in motor learning. We discovered that the ChC is essential for improving the direction-tuning of targeted cells by suppressing irrelevant activity at the population level. ChC shaped this sophisticated inhibitory motif among a specific set of neuron types. Thus, ChCs primarily take a “select and focus” strategy rather than serve as a uniform gain setter.

## Results

### Improved directional control of movement on multi-textured ball maze

Previous studies have shown that ChCs increased activity during locomotion ^17, 18^. The premotor cortex (M2, anteromedial agranular cortex, homologous to supplementary motor areas in the primate brain) is involved in gating sensory inputs and motor outputs for movement planning, fine action control, and decision-making ^19–21^. To investigate the role of ChCs in controlling the activity of M2 neurons during locomotion, we used a spatial navigation task that requires goal-directed motor control ^22^. Mice were trained to traverse a multi-textured ball maze with four distinctive tactile surface cues and to reach a goal spot on the ball, where a water reward was given (**Fig. 1a**, See Methods). The task required directional motor control accurately aimed to the goal spot. While animals were navigating the ball maze, we recorded neuronal activity of the L2/3 premotor population using Ca^2+^ imaging via two-photon microscopy. Mice with water-restriction (experimental group) showed reward-seeking behavior on the ball maze, developed a preferred pathway, and generated goal-oriented locomotion with learning whereas mice *ad libitum* water (control group) did not (**Fig. 1b**). The number of successfully reaching the goal spot increased in the experimental group but not in the control group (**Fig. 1e**). Time spent in the goal (rewarded) quadrant (Q1) increased with training (**Fig. 1f**). We further examined the relationship between motor control and reward-seeking behavioral performance by analyzing the parameters of motor control. To examine whether general movement parameters such as speed and acceleration could be the main cause for these improvements, we compared the motor control between experimental and control conditions. The movement of the control group showed similar general motor parameters (e.g., speed and acceleration) to the experimental group (**Fig. 1c, d**). Overall latency to the reward decreased with training, indicating the increased effectiveness of the animals’ movements at achieving rewards (**Fig. 1g**). We observed that mice showed increased proximity to the goal with learning, by tuning their movement to the goal (**Fig. 1h**). The movement accuracy as the percentage of goal-aimed movements increased with training (**Fig. 1i**), indicating that the enhanced movement direction (MD) controls toward the goal spot. Mice in the experimental group showed better control in turning toward the goal during navigation than the control (**Fig. 1j, k**). These results indicate that the directional control of movement is improved with learning.

**Figure 1.**
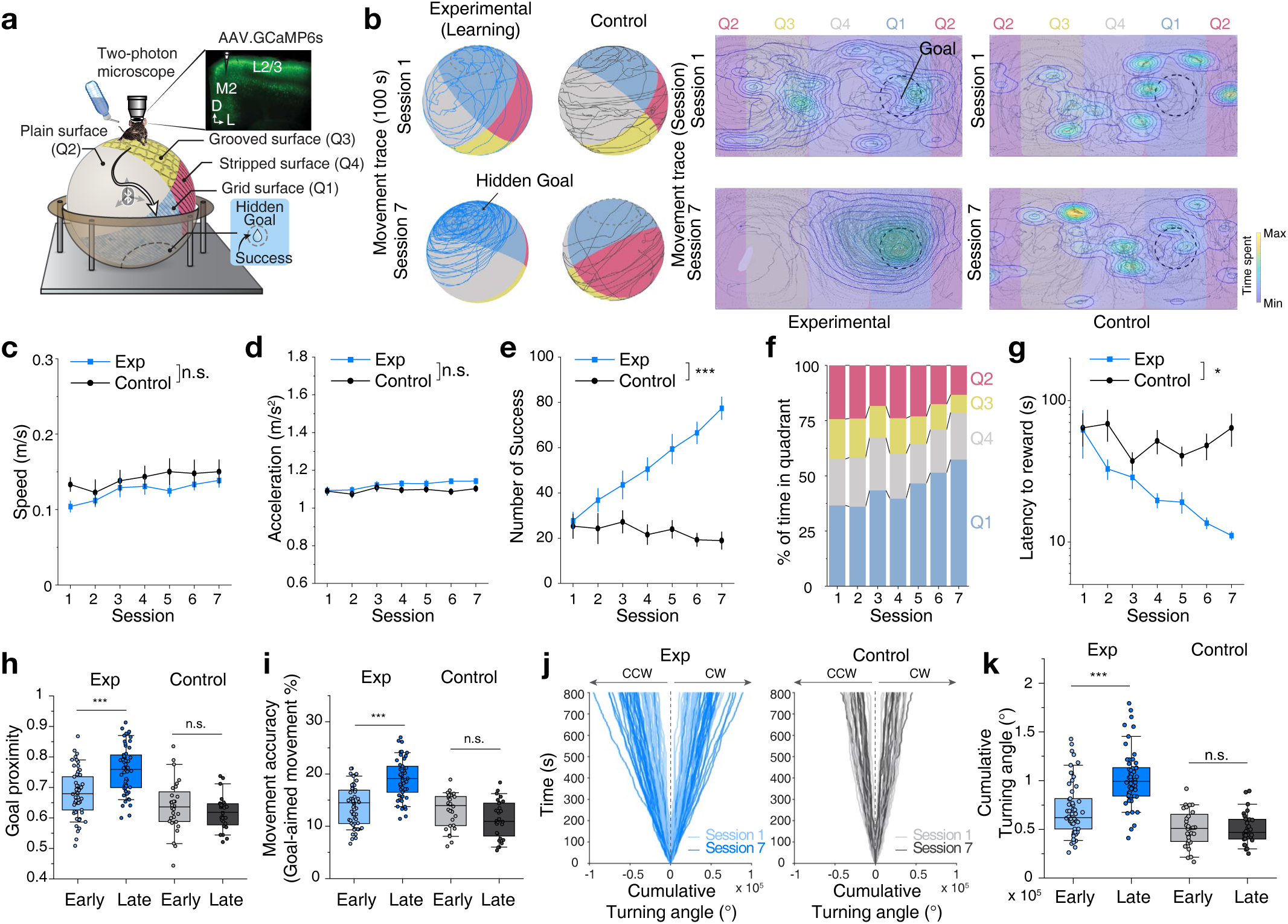
Improved directional control of movement with learning. **(a)** Schematic of the experimental setup for the spatial navigation task on a multi-textured floating ball maze, in which head-fixed mice freely navigate toward a goal spot in the target quadrant (Q1) based on tactile cues. Two-photon calcium imaging was simultaneously performed to monitor neuronal activity in layer 2/3 of premotor cortex (M2). **(b)** Representative movement trace (100 s) of mice in the experimental (water restricted) and control (water *ad libitum*) groups navigating on the multi-textured floating ball maze from training sessions 1 to 7 (left) and their 2-dimensional projection (right). A circle with a dashed line indicates the goal spot on the ball. Heat map and contour lines of the times of mice spent on the location are presented. **(c)** Average movement speed (n = 27 for the experimental group; n = 14 for the control group; Two-way repeated measures ANOVA, *F_group_* = 3.85, *P* = 0.072). **(d)** Average movement acceleration (two-way repeated measures ANOVA, *F_group_* = 154, *P* = 0.24). **(e)** Average number of successes obtained in training sessions (two-way repeated measures ANOVA, *F_group_* = 26.97, *P* = 1.73 × 10^−4^). **(f)** Percentage of time spent in each quadrant during navigation in the experimental group. **(g)** Average latency to reward (two-way repeated measures ANOVA, *F_group_* = 7.43, *P* = 0.017). **(h)** Average goal proximity (two-tailed *t*-test, *t* = −5.19, *P* = 1.03 × 10^−6^ for the experimental group; *t* = 1.15, *P* = 0.26 for the control group). **(i)** Movement accuracy (two-tailed *t*-test, *t* = −7.04, *P* = 2.01 × 10^−10^ for the experimental group; *t* = 1.74, *P* = 0.09 for the control group). **(j)** Cumulative turning angle over time in Sessions 1 and 7. **(k)** Comparison of cumulative turning angle between the experimental and control groups (two-tailed *t*-test, *t* = −5.84, *P* = 5.76 × 10^−8^ for the experimental group; *t* = 0.019, *P* = 0.98 for the control group). n.s., not significant; \*\**P* < 0.01; \*\*\**P* < 0.001; Error bars indicate s.e.m. In the box plot, the midline, box size, and whisker indicate median, 25-75th percentile, and 10-90th percentile, respectively.

### Motor learning refines direction coding through suppression of irrelevant activity

To determine how neurons mediate precise motor control with learning, we examined direction coding of M2 neurons at the cellular and population levels. To test whether M2 is required for neuronal adaptation to control task-related motor behavior, we used an adeno-associated virus (AAV)-based approach to express tetanus toxin light chain (TeTxLC), which abolishes presynaptic vesicle release ^23, 24^, in M2 excitatory neurons (**Extended Data Fig. 1a**). Mice expressing TeTxLC in M2 excitatory neurons (CaMKII-TeTxLC mice) showed deficits in learning the task (**Extended Data Fig. 1e**). The reward-seeking behavioral parameters did not improve with training (**Extended Data Fig. 1e-j**), suggesting that M2 neurons play an essential role in mediating the motor learning required for the task.

We performed two-photon Ca^2+^ imaging in L2/3 of M2 with a genetically encoded calcium indicator, GCaMP6s while mice navigated on the ball maze (**Extended Data Fig. 2a**). Consistent with the established role of M2 in motor planning, overall population activity in L2/3 correlated with locomotion (**Extended Data Fig. 2b**). Individual premotor neuron activities bi-directionally changed in a time-locked fashion at movement onset (**Extended Data Fig. 2c**). The majority of neurons were movement-related (total 98.0%; positively correlated: 79.7 ± 3.3%; negatively correlated: 18.4 ± 3.2%; and not correlated: 2.0 ± 0.4%; see Methods for classification), and the proportion of movement-related neurons in the population remained relatively constant throughout training sessions (**Extended Data Fig. 2d**).

We next measured direction selectivity of individual premotor neurons over training and analyzed the changes. Distinct neuronal populations responded to different MDs (MDs) (**Fig. 2a, Extended Data Fig. 2e-g**). A subset of L2/3 neurons exhibited tuning to the MD which was defined as its preferred direction (PD). Its topological map was arranged in a salt-and-pepper pattern in the M2 area (**Extended Data Fig. 2i, j**).

**Figure 2.**
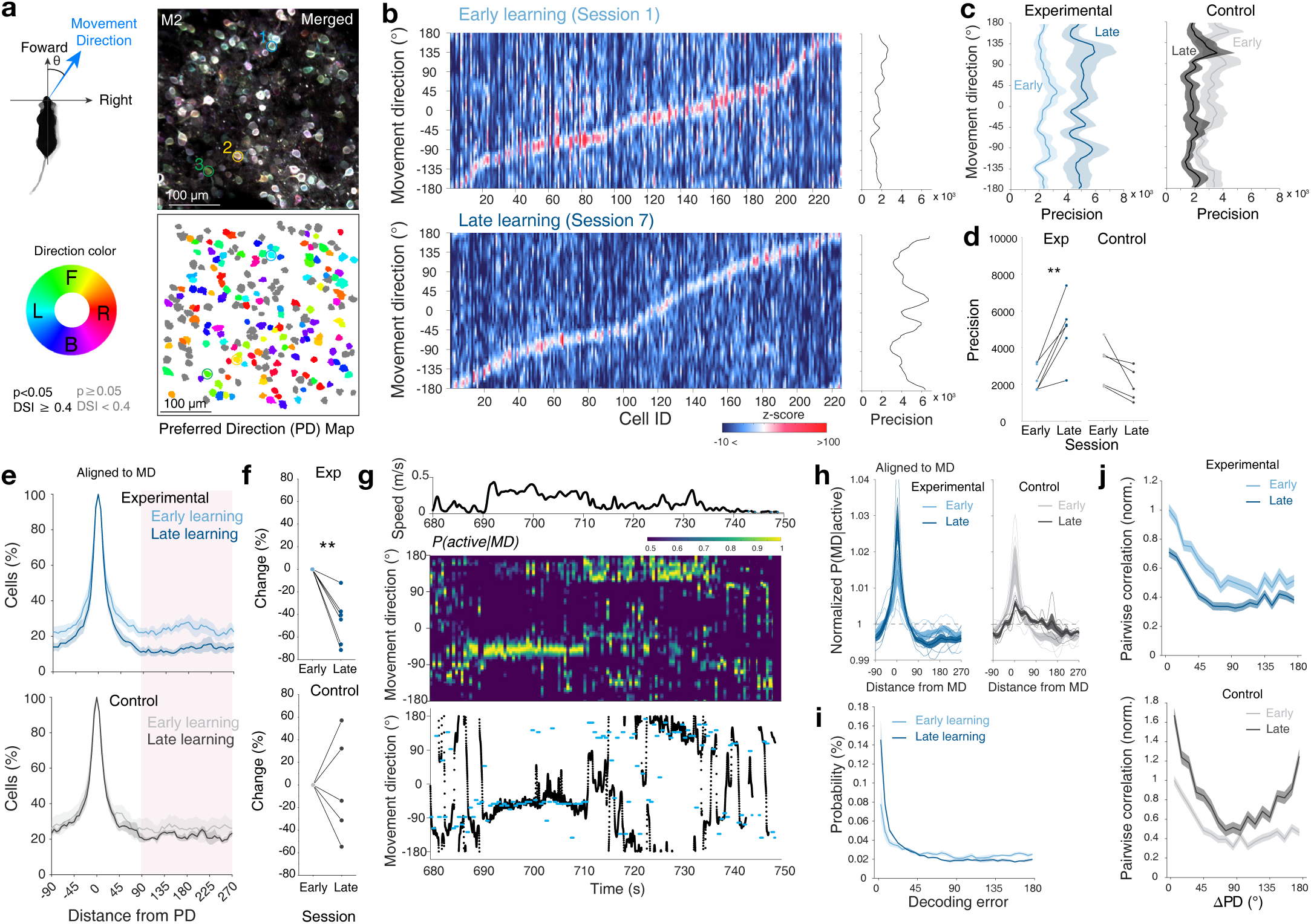
Motor learning refines direction coding through the suppression of irrelevant activity. **(a)** Schematic of movement direction based on forward-backward and right-left speeds. An average pixel-based activity map of movement direction (top). Corresponding merged activity map (top). Color-coded map of preferred directions of direction-tuned neurons (von Mises fitting *P* < 0.05 and Direction Selectivity Index, DSI ≥ 0.4), colored according to their preferred direction. Non direction-tuned neurons (DSI < 0.4) are shown in gray. **(b)** Example heatmap of premotor neurons’ tuning for movement direction in early learning (session 1, top) and late learning (session 7, bottom) from a mouse in the experimental group. Data is sorted from the location of peak likelihood probability P (active|MD). Corresponding precisions of population responses for each movement direction (right). **(c)** Changes of average precision curves of population responses in the experimental and control groups. **(d)** Comparison of population response precisions between early and late sessions (n = 6 mice for the experimental group, two-tailed paired t-test, *t* = −4.98, *P = 0.0042*; n = 5 mice for the control group, *t* = 2.56, *P = 0.063*). **(e)** Normalized percentage of active cells in the population as a function of distance from the preferred direction (n = 830 cells for early, n = 937 cells for late in the experimental group; n = 705 cells for early, n = 671 cells for late in the control group). **(f)** Changes in the percentage of active cells from the early session and late session (two-tailed Student’s t-test. *t* = 5.149, *P* = 0.0036 for the experimental group; *t* = 0.106, *P* = 0.920 for the control group). **(g)** Movement speed (top), posterior probabilities (middle), and corresponding actual and decoded movement direction estimated with maximum a posteriori (MAP)(bottom). **(h)** Changes in posterior probabilities normalized by chance level as a function of distance from movement direction with learning for experimental and control groups. **(i)** Probability distributions of decoding error. **(j)** Pairwise correlations with respect to ΔPD normalized by early session Session 1 (angular difference in preferred direction between neuronal pairs, n = 58456 pairs for Session 1, n = 67143 pairs for Session 7 for the experimental group; n = 51285 pairs for Session 1, n = 45156 pairs for Session 7 for the control group). \*\**P* < 0.01; Error bars and shading indicate s.e.m.

We estimated the likelihood distributions of neural tuning curves as a function of MD (**Fig. 2b**). Premotor neurons showed the highest likelihood to fire at their PD. We defined population tuning precision as the reciprocal of the variance of total neural activity given MD. Tuning of premotor neurons to MD became more precise over sessions. The precision of population responses increased for all MDs in the experimental group, but not in the control group (**Fig. 2c-d**). We next estimated the probability of the cell being active as a function of the distance between MD and the cell’s PD. When the MD was far from the cell’s PD (distance from PD > 90°), the probability of being active decreased with learning (**Fig. 2e**). When the MD was opposite to the cell’s PD, the probability of the cell being active was significantly lower in Session 7 than in Session 1 for mice in the experimental condition, but not in the control condition (**Fig. 2f**).

We took a Bayesian approach to decode the MD from neural activity (See Methods). We computed the posterior probability density function to provide the most likely estimate of the mouse’s MD during navigation (**Fig. 2g, Extended Data Fig. 3a-c**). Moment-to-moment MDs were decoded by the estimated MD corresponding to the maximum value of the posterior probability density function. During behavior, posterior probability density functions of MD sharpened over sessions in the experimental group, enhancing near the MD and waning at null MDs (**Fig. 2h**). In contrast, posterior functions were flattened over sessions in the control group. Consequently, we observed lower decoding errors in the experimental group (**Fig. 2i**). Similarly, we tested neural coding of MD based on a population vector (PoV) as a vector sum of PDs of a sparse population of active direction-tuned neurons at a given moment ^25, 26^ (**Extended Data Fig. 3d**). In this coding scheme, MDs were also flexibly encoded in a moment-to-moment manner from the distinct sparsely distributed activity of neuronal ensembles (**Extended Data Fig. 3e, f**). The fraction of decoded direction (PoV around 0°) increased and that of anti-decoded direction (PoV around 180°) decreased in Session 7 in the experimental group, but not in the control group, indicating an increased coding accuracy associated with learning (**Extended Data Fig. 3g-i**). We further measured angular errors between the active neuron’s PD and MD to see how directional neuronal activity was accurately aligned with MD on a moment-to-moment basis. Similar to the change in decoding error, the fraction of PD increased and that of anti-PD decreased in Session 7 in the experimental mice, whereas the fraction of anti-PD increased in Session 7 in the control mice, indicating that the fraction of irrelevant activity reduced with learning (**Extended Data Fig. 3j, k**). To test whether the motor learning mediates a direction-dependent change in network connectivity, we compared pairwise correlations between neuronal pairs of diametrically opposed PDs. The pairwise correlations of neuronal pairs decreased in the experimental group as a function of the difference in their PDs, whereas they increased over sessions in the control group (**Fig. 2j**). These results suggest that the observed reduction of incorrectly tuned responses shapes the direction coding of the M2 population.

### Manipulation of parvalbumin interneurons disrupts global neural activity

To determine how neuronal ensembles suppress irrelevant activity during learning, we asked how interneurons might be uniquely involved. Several GABAergic interneuron subtypes have been hypothesized to serve specific roles in regulating cortical functions by forming circuit motifs with features like recurrence and feedforward inhibition ^27^. We tested the regulative role of two major interneuron groups that are known to form inhibitory synapses at specific compartments of excitatory PyNs and comprise 70% of all cortical interneurons ^28^; Specifically, PV-INs inhibit perisomatic regions and SOM-INs target distal dendrites ^29^. We bilaterally silenced activity of PV-INs or SOM-INs during behavior by expressing an inhibitory opsin, eNpHR3.0, in the M2. We found that inactivation of premotor PV-INs disrupted goal-directed motor performance, but inactivation of SOM-INs did not, suggesting that perisomatic inhibition mediated by PV-INs may be involved in circuit motifs that guide task-related directional control of movement (**Fig. 3a, Extended Data Fig. 4**).

**Figure 3.**
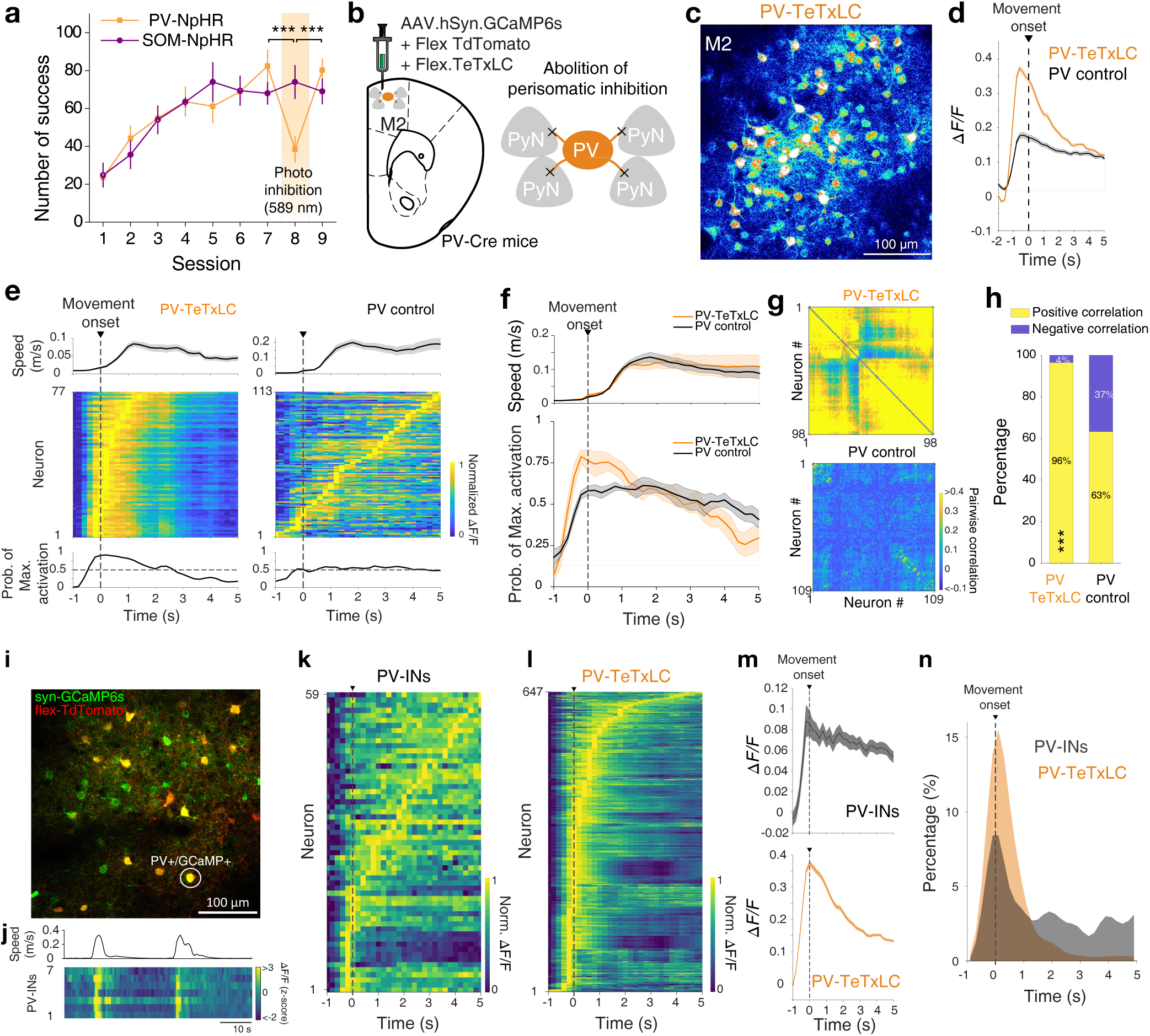
Manipulation of parvalbumin interneurons alters global neural activity. **(a)** Behavioral performance of the number of successes with training and photo-inhibition. Inactivation of PV-INs, but not SOM-INs, resulted in a deficit in finding a hidden goal (n = 8 mice for PV-NpHR, one-way repeated measures ANOVA, *F_session_*= 30.0, *P* = 8.69 × 10^−6^, Fisher multiple comparisons tests, \*\*\**P* <0.001; n = 5 mice for SOM-NpHR, *F_session_* = 0.42, *P* = 0.67). **(b)** Schematic of blockage of GABA release from PV-INs by expressing tetanus toxin light chain (TeTxLC) in PV-Cre mice. **(c)** Representative field of view showing increased excitability of premotor neurons in PV-TeTxLC mice. **(d)** Average fluorescence transient traces of premotor neurons aligned to movement onset in PV-TeTxLC and PV control mice (n = 647 cells from 6 mice for PV-TeTxLC, n = 480 cells from 7 mice for PV control). **(e)** An example of movement speed (top), the normalized activity of neurons (middle), and probability of maximum neuron activation (bottom) aligned to movement onsets in PV-TeTxLC and PV control. **(f)** Average movement speed (top) and corresponding changes of the probability of maximum neuron activation (bottom) aligned to movement onsets. **(g)** Example pairwise correlation matrices of PV-TeTxLC and PV control mice. **(h)** Population fraction of neuronal pairs with positive and negative correlation (n = 31660 pairs for PV-TeTxLC, n = 24061 pairs for control; chi-square test, *χ*^2^ = 1.03 x 10^4^, *P* = 0) **(i-n)** The activity of PV-INs and ChCs related to the movement initiation. **(i)** An example field of view showing premotor neurons expressing GCaMP6s (green) and FLEX-tdTomato (red) in PV-Cre mice. **(j)** An example trace of movement speed (top) and corresponding heatmap of fluorescent transient signals of PV-INs (bottom). **(k)** The normalized activity of PV-INs aligned to movement onset (n = 59 cells from 7 mice in PV control). **(l)** The normalized activity of neurons aligned to movement onset in PV-TeTxLC group (n = 647 cells from 6 mice in PV-TeTxLC). **(m)** Average ΔF/F traces of PV-INs (top) and neurons in PV-TeTxLC group (bottom) aligned to movement onset. **(n)** Probability distributions of peak activity timing of PV-INs and neurons in PV-TeTxLC group aligned to movement onset. \*\*\**P* < 0.001; Error bars and shadings indicate s.e.m.

Perisomatic inhibition from PV-INs provides powerful inhibitory control over the principal neuron population ^29–32^. To further examine the role of PV-IN perisomatic inhibition in local neuronal activity during motor control, we abolished presynaptic GABA release from PV-INs by expressing Cre-dependent TeTxLC virus in PV-Cre mice (PV-TeTxLC mice) and monitored the M2 neuron activity (**Fig. 3b**). Consistent with the notion that PV-INs are critical regulators of cortical excitatory and inhibitory (E/I) balance ^33, 34^, the abolishment of perisomatic inhibition in PV-TeTxLC mice led to excessively increased overall activity of M2 neurons during movement (**Fig. 3c-d**) ^35^ and hypersynchronization at movement initiation, which disrupted the propagation of sequential network activity and decreased the sparseness of neural activity during subsequent movements (**Fig. 3e, f**). Accordingly, the pairwise correlation and the proportion of pairs with positive correlations were significantly higher in PV-TeTxLC mice than in PV control mice (**Fig. 3g-h**). A significant proportion of PV-INs also showed increased activity for movement initiation (**Fig. 3j**), suggesting that the excessive activity of M2 neurons in PV-TeTxLC mice may result from unmasked responses of neurons that were previously inhibited by PV-INs (**Fig. 3k-n**). These results suggest that PV-IN perisomatic inhibition is involved in task-related motor control, but this manipulation also leads to global excitability changes that limits the applicability of current approaches to specify mechanisms of directional motor control.

### Sparse, sequential activity is preserved in ChC manipulation

In our attempt to specify the underlying mechanisms of perisomatic inhibition, we targeted ChCs by using the ChC-specific transgenic mouse line (Nkx2.1-2a-CreER::Ai14 as ChC control mice, Nkx2.1-2a-CreER::Flex-FlpO mice as ChC-Flp mice, and Vipr2-Cre) ^36^. This genetic strategy allowed us to selectively label ChCs, including their distinctive axonal arborizations and cartridge terminals (**Extended Data Fig. 5a-c**). A larger proportion of ChCs were found in the upper L2/3 of M2, with a smaller number present in L5 (mean soma depth ± s.e.m = 178.9 ± 10.7 μm from pial surface) (**Extended Data Fig. 5d**). We confirmed that the axonal cartridges of ChCs made synapses on the AISs of neighboring neurons (**Fig. 4a**). To find out if ChC manipulation impacts global network excitability, we abolished presynaptic inhibitory GABAergic inputs to the AISs of neighboring neurons by expressing FlpO-dependent TeTxLC (AAV9-CAG-dfrt-TeTxLC-HA-WPRE) bilaterally in ChC-Flp mice and monitored activity of M2 neurons (**Fig. 4b**). In contrast to the hypersynchronous neuronal activity in PV-TeTxLC mice, we did not observe a global change. Instead, we observed sparse, sequential activity propagation in both ChC-TeTxLC and control mice during movement (**Fig. 4c, d**). Pairwise correlation indicated local, not global, correlated activity between neuronal pairs in both groups (**Fig. 4e**); The proportions of pairs with positive and negative correlations were similar between the two groups (**Fig. 4f**). In contrast to PV-INs, a majority of ChCs showed increased activity during movement rather than movement initiation (**Fig. 4g**), and ChC-TeTxLC manipulation did not evoke the excessive activity of M2 neurons at movement onset shown in PV-TeTxLC mice (**Fig. 4h-k**). These results demonstrate that ChCs have a distinct activity profile from PV-INs and that overall network excitability is not globally altered by silencing the activity of ChCs.

**Figure 4.**
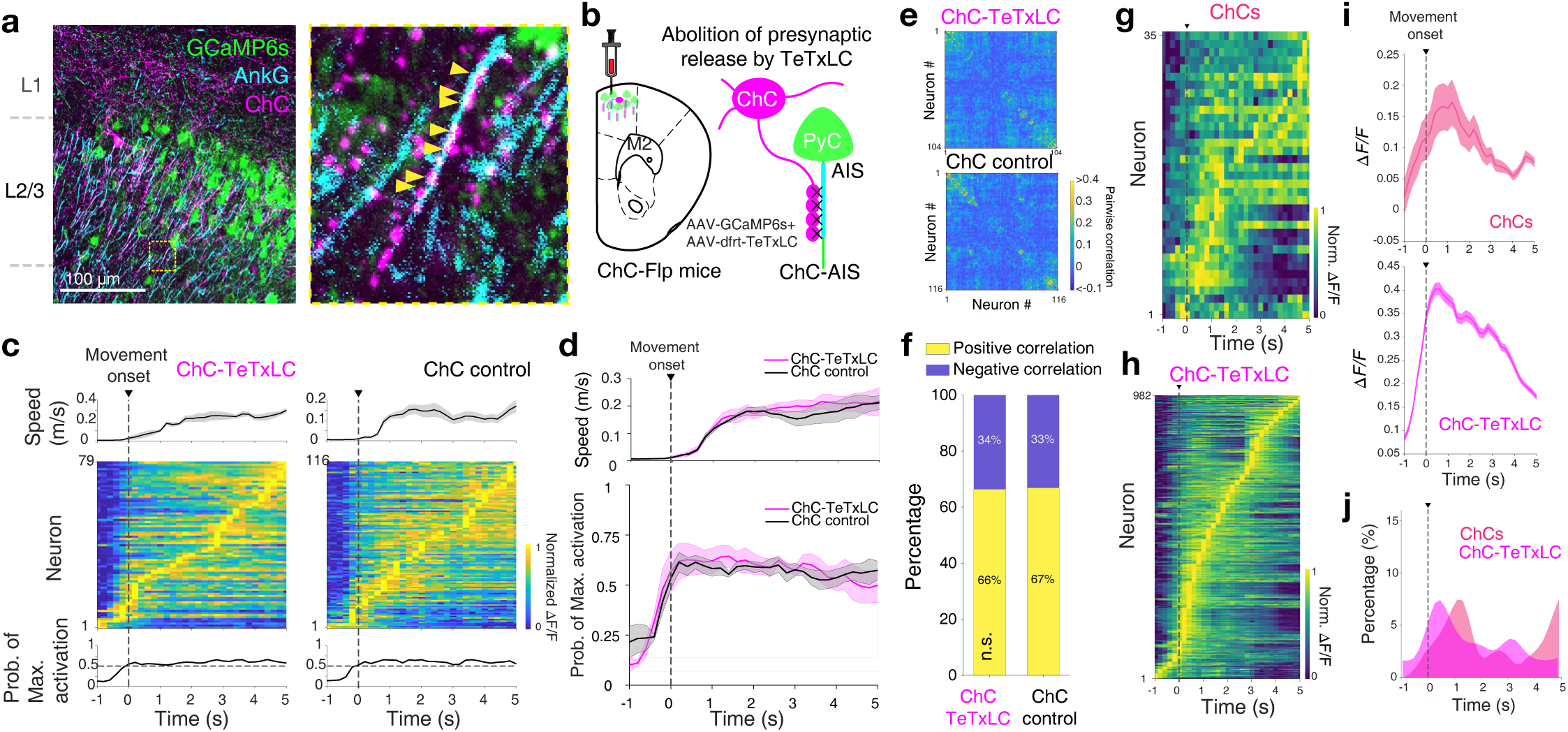
Sparse, sequential activity is preserved in ChC manipulation. **(a)** Example image of the axonal projection of ChC (expressing mCherry, in magenta) to AISs (post-hoc AnkG staining, in cyan) of neighboring L2/3 neurons (expressing GCaMP6s, in green). Axonal boutons of ChC innervating an AIS (right). **(b)** Schematic of selective abolition of GABA release from ChCs in M2 by expressing tetanus toxin light chain (TeTxLC) in ChC-Flp mice (Nkx2.1-2a-CreER::Flex-FlpO mice). **(c)** An example of movement speed (top), the normalized activity of neurons (middle), and probability of maximum neuron activation (bottom) aligned to movement onsets in ChC-TeTxLC and ChC control. **(d)** Average movement speed (top) and corresponding changes in the probability of maximum neuron activation (bottom) aligned to movement onsets. **(e)** Example pairwise correlation matrices of ChC-TeTxLC and ChC control mice. **(f)** Population fraction of neuronal pairs with positive and negative correlation (n = 14082 pairs for ChC-TeTxLC, n = 57956 pairs for ChC control; chi-square test, *χ*^2^ = 1.21, *P* = 0.27). **(g-j)** The activity of ChCs related to the movement initiation. **(g)** The normalized activity of ChCs aligned to movement onset (35 cells from 3 mice). **(h)** The normalized activity of neurons aligned to movement onset in ChC-TeTxLC group (n = 982 cells from 9 mice). **(i)** Average ΔF/F traces of ChCs (top) and neurons in ChC-TeTxLC group (bottom) aligned to movement onset. **(j)** Probability distributions of peak activity timing of ChCs and neurons in ChC-TeTxLC group aligned to movement onset. Error bars and shading indicate s.e.m.

### Chemogenetic inhibition of ChCs disrupts the direction selectivity of premotor neurons

The TeTxLC experiments suggest that the major role of ChCs is not control of general network gain, but more target-specific control of neighboring neurons. We tested whether ChC suppression would affect the direction tuning of M2 neurons during the behavioral task. We bilaterally expressed FlpO-dependent chemogenetic silencer (AAV9-CAG-dfrt-hM4Di-mCherry) in the M2 of ChC-Flp mice to suppress the activity of ChCs upon the application of clozapine N-oxide (CNO), an hM4Di agonist (**Fig. 5a**). We first verified the effectiveness of CNO in brain slices. Whole-cell electrophysiological recordings showed that CNO application reduced the activity of ChCs expressing hM4Di (**Fig. 5b**). After 7 days of training, we administered either saline or CNO. CNO administration in the ChC-Flp:hM4Di mice significantly impaired goal-directed navigation performance and turning movement (**Fig. 5c-f**). CNO or saline injection alone had no effect on behavioral performance (ChC-Flp:hM4Di mice with saline, ChC-Flp mice with saline or CNO injection) (**Fig. 5d**).

**Figure 5.**
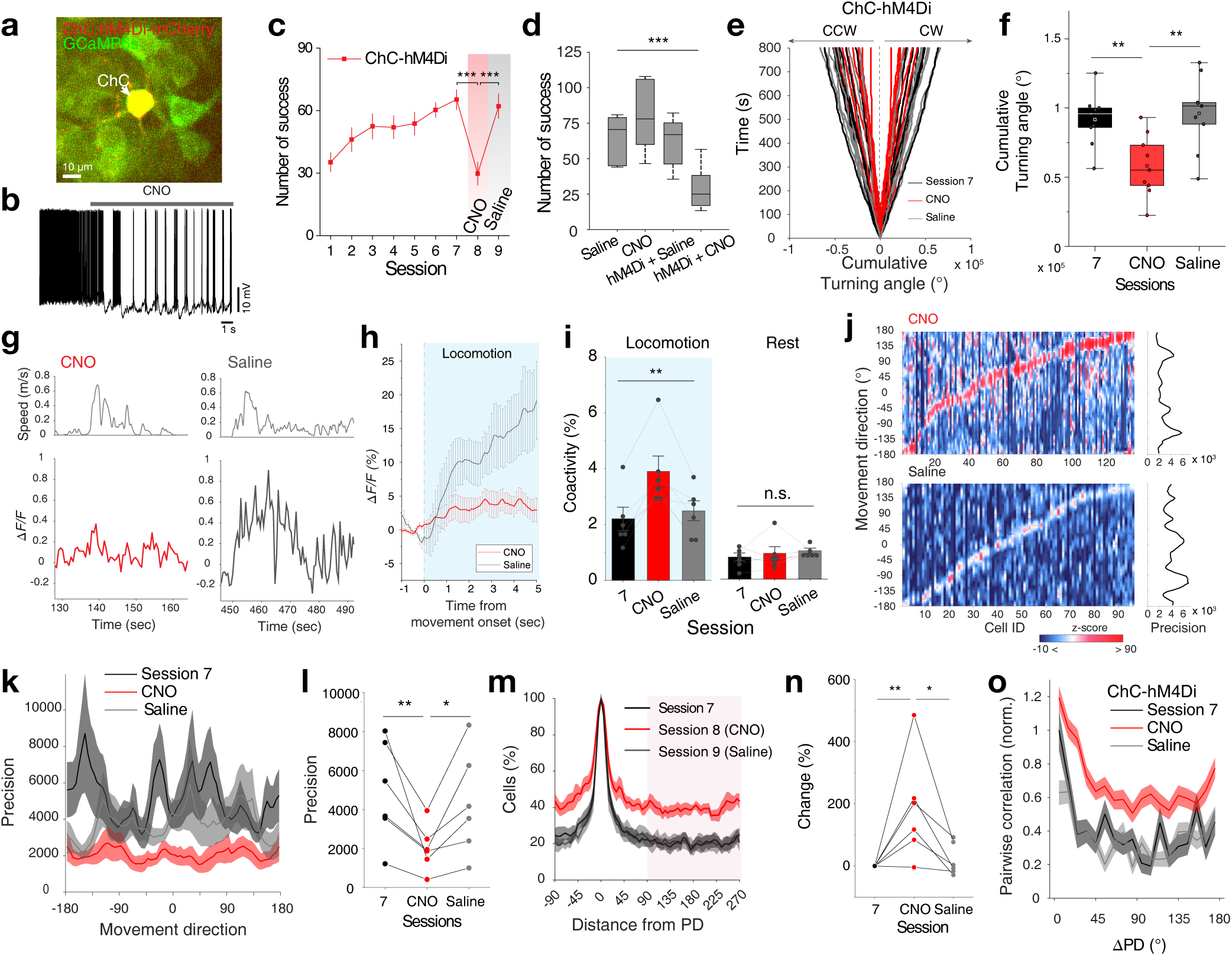
Chemogenetic inhibition of ChCs disrupts the direction selectivity of premotor neurons. **(a)** *In vivo* two-photon Ca^2+^ imaging of ChC and neighboring neurons expressing GCaMP6s and selectively coexpressing the chemogenetic silencer, hM4Di-mCherry in ChCs of M2, in a FlpO-dependent manner, using Nkx2.1-2a-CreER::Flex-FlpO mice (ChC-Flp mice). **(b)** Bath-application of CNO (10 μM) reduced the firing rate in ChCs expressing hM4Di. Example whole-cell current-clamp recording from a ChC. **(c)** The average number of successes of ChC-hM4Di mice increased with training from sessions 1 to 7. CNO or saline injection was followed on session 8 or 9 (n = 9 mice, one-way repeated measures ANOVA, *F_session_* = 31.1, *P* = 3.09 × 10^−6^; Fisher multiple comparisons tests, Session 7 vs. CNO, *P* = 2.40 × 10^−6^; Session CNO vs. Saline, *P* = 7.58 × 10^−6^; Session 7 vs. Saline, *P* = 0.53). **(d)** Cumulative turning angle of ChC-hM4Di mice across conditions (Session 7, CNO, and Saline). **(e)** Comparison of cumulative turning angle across conditions (one-way repeated measures ANOVA, *F_session_* = 8.99, *P* = 0.0024; Fisher multiple comparisons tests, Session 7 vs. CNO, *P* = 0.0035; Session 7 vs. Saline, *P* = 0.45; CNO vs. Saline, *P* = 0.0013). **(f)** Comparison of performance between conditions (n = 6 mice for ChC control with saline or CNO; n = 9 mice for ChC-hM4Di with saline or CNO, one-way ANOVA *F_sesion_* = 9.50, *P* = 2.06 × 10^−4^). **(g)** Example ΔF/F traces of a ChC during locomotion in CNO and saline conditions. **(h)** Average ΔF/F traces of ChCs aligned to movement onset in CNO and saline conditions (6 ChC-hM4Di mice, n = 497 cells for CNO; n = 475 cells for Saline). **(i)** Coactivity percentage of neurons during periods of locomotion and rest. Friedman test, *χ*^2^ = 9.33, *P* = 0.0094 for locomotion; *χ*^2^ = 0.33, *P* = 0.85 for rest). **(j)** Example tuning curves of each individual premotor neuron for movement direction in CNO (top) and saline (bottom) conditions. Data is sorted from the location of peak likelihood probability P (active|MD). Corresponding precisions of population responses for each movement direction (right). **(k)** Average precision curves of population responses across later learning, CNO, and saline conditions. **(l)** Comparison of population response precisions (one-way repeated measures ANOVA, *F_session_* = 6.39, *P* = 0.016; Fisher multiple comparisons tests, Session 7 vs. CNO, *P* = 0.0068; Session CNO vs. Saline, *P* = 0.023; Session 7 vs. Saline, *P* = 0.486). **(m)** Normalized percentage of active cells in the population as a function of distance from the preferred direction (n = 484 cells for Session 7, n = 407 cells for CNO, n = 582 cells for Saline). **(n)** Changes in the percentage of active cells across conditions. (one-way repeated measures ANOVA with Greenhouse-Geisser correction, *F_session_* = 7.26, *P* = 0.037; Fisher multiple comparisons tests, Session 7 vs. CNO, *P* = 0.006; Session CNO vs. Saline, *P* = 0.011; Session 7 vs. Saline, *P* = 0.372). **(o)** Pairwise correlations with respect to ΔPD normalized by Session 7 (angular difference in preferred direction between neuronal pairs, n = 17933 pairs for Session 7; n = 24205 pairs for CNO; n = 22735 pairs for Saline). \**P* < 0.05, \*\**P* < 0.01, \*\*\**P* < 0.001; n.s., not significant. Error bars and shading indicate s.e.m.

To determine the role of ChC activity in the direction tuning of M2 neurons, we co-expressed AAV9-CAG-dfrt-hM4Di-mCherry and AAV1-hSyn-GCaMP6s into the L2/3 M2. We then used chemogenetic manipulations and performed Ca^2+^ imaging of ChCs and their neighboring neurons during the behavioral task. In line with the *ex vivo* results, ChC activity was suppressed upon CNO administration during locomotion (**Fig. 5g, h**). Consequently, we observed that the coactivity of M2 neurons increased upon CNO application during locomotion but not during rest (**Fig. 5i**), consistent with recent work demonstrating the inhibitory postsynaptic effects of ChCs on M2 neurons *in vivo* ^17, 31^. We compared the likelihood distributions of neural tuning curves as a function of MD with and without CNO administration. We observed that neurons are more likely to be active at non-PDs in the CNO (ChC-silenced) condition than in the saline condition (**Fig. 5j**), resulting in less precise population responses to the MD (**Fig. 5k, l**). The probability of the cell being active at a certain MD increased upon CNO administration (**Fig. 5m**). When the MD was at the opposite of the cell’s PD, the probability of the cell being active was higher after CNO administration than it had been in Session 7 or in the saline condition (**Fig. 5n**). CNO application increased pairwise correlations between neuronal pairs as opposed to the enhanced decorrelation between pairs seen with learning (**Fig. 5o**). These results suggest that the suppression of ChC activity increased behaviorally irrelevant neural activity, impairing the precise coding of movement.

### Variability of ChC activity increases during learning

To examine the activity dynamics of ChCs during learning, we expressed GCaMP6s in the L2/3 ChCs and recorded ChC activity while mice performed the behavioral task (**Fig. 6a**). Consistent with previous reports ^17, 18^, ChCs displayed increased activity during episodes of locomotion (**Fig. 6b**), with a more synchronized pattern of activity in Session 1. In Session 7, activity patterns became more diverse and desynchronous, which led to the formation of subclusters of ChCs (**Fig. 6c**). To quantify this shift, we calculated the pairwise correlations between ChCs during the behavioral task and found that the correlation decreased and ChCs formed subclusters as learning progressed (**Fig. 6d, e**). We next examined how temporal relationships between ChCs and M2 neurons change with learning. Various temporal relationships in activity were shown between ChCs and neighboring M2 neurons (**Fig. 6f**). We calculated cell-to-cell activity correlations (Pearsons’ correlation) in pairs of ChCs and M2 neurons during locomotion and rest epochs and compared their distributions of correlation coefficients. In Session 1, the proportion of high correlation coefficient (corr(r) > 0.6) was decreased during locomotion compared to rest, reflecting the increased decorrelation. In Session 7, the overall proportions of positive correlation coefficient (corr(r) > 0) and of negative coefficient (corr(r) < 0) were remarkably decreased during locomotion compared to rest, indicating the enhanced decorrelation in gross pairs of ChC and M2 neuron (**Fig. 6g**). These results suggest that ChC activity becomes more heterogeneous with learning and such decorrelated temporal relationship between ChCs and neighboring M2 neurons implies an increased specificity of population activity control.

**Figure 6.**
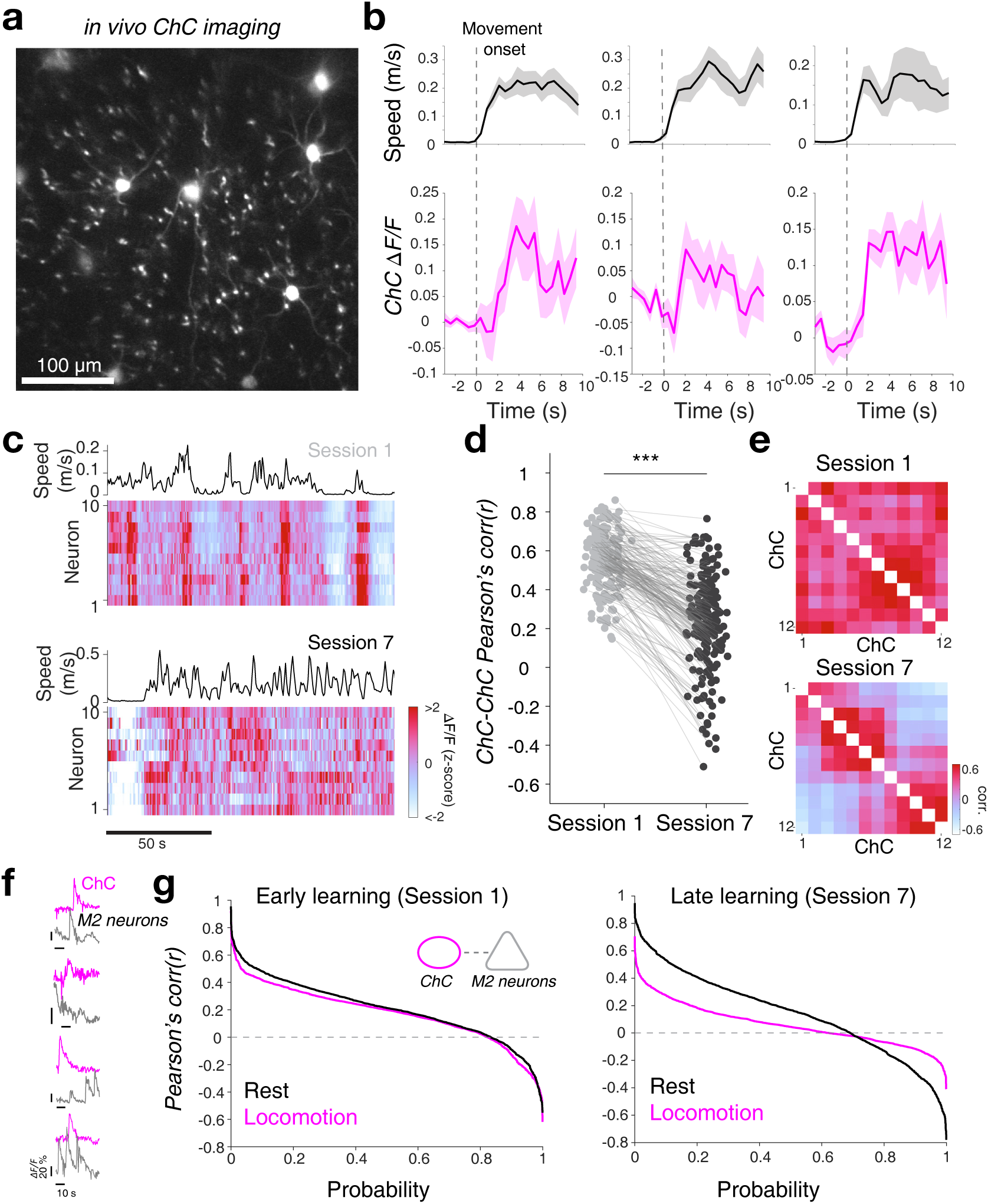
Variability of ChC activity increases during learning. **(a)** Representative field of view showing L2 ChCs expressing GCaMP6. **(b)** Example fluorescence traces from ChCs that show increased activity during episodes of locomotion. **(c)** Movement speed and the heatmap of ChC activity during navigation in Sessions 1 and 7. The synchronous activation of ChCs in Session 1 during locomotion becomes asynchronous in Session 7. **(d)** Changes in the correlation between a pair of ChCs (n = 189 pairs in 35 cells from 3 mice; two-tailed Wilcoxon signed-rank test, *Z* = 11.92, *P* = 0). **(e)** Example correlation matrix of ChCs in Sessions 1 and 7. **(f)** Various temporal relationships between activity of ChC (magenta) and neighboring M2 neurons (gray) during episodes of locomotion in ChC-hM4Di mice. **(g)** Cell-to-cell Pearson’s correlation of ChC-M2 neuron pairs in Sessions 1 and 7 during locomotion and rest epochs (4 ChC-hM4Di mice, n = 1957 pairs for Session 1, two-sample Kolmogorov-Smirnov test, locomotion vs. rest, *D* = 0.076, *P* = 2.12 × 10^−5^; n = 1723 pairs for Session 7, locomotion vs. rest, *D* = 0.28, *P* = 4.049 × 10^−61^). \*\*\**P* < 0.001; Shadings indicate s.e.m.

### Abolition of ChC inhibition impairs refinement of direction coding and motor learning

We asked if the abolition of ChC-mediated inhibition would affect the refinement of direction coding and learning ability. We abolished presynaptic ChC GABA release by expressing FlpO-dependent TeTxLC (AAV9-CAG-dfrt-TeTxLC-HA-WPRE) in ChC-Flp mice, and performed two-photon calcium imaging in M2 neurons across sessions (**Fig. 7a-c**). ChC-TeTxLC mice showed a deficit in learning the task, indicating the necessary role of axo-axonic inhibition for learning goal-directed motor control (**Fig. 7e-l**). Given the vast difference in the numbers of PV-INs and ChCs ^37^, it is possible that the suppression of a similar number of PV-INs has similar effects to the effects exerted by the selective suppression of ChCs. To test this possibility, we abolished the inhibition presynaptic GABA release of a fraction of PV-INs that corresponds approximately to the percentage of ChCs in the premotor network (**Extended Data Fig. 6a-c**). The sparse, sequential activity was observed in sparse PV-TeTxLC mice similar to PV control (**Extended Data Fig. 6d**). However, in contrast to ChC-TeTxLC mice, the sparse PV-TeTxLC mice showed normal improvement of learned performance (**Extended Data Fig. 6e, f**).

**Figure 7.**
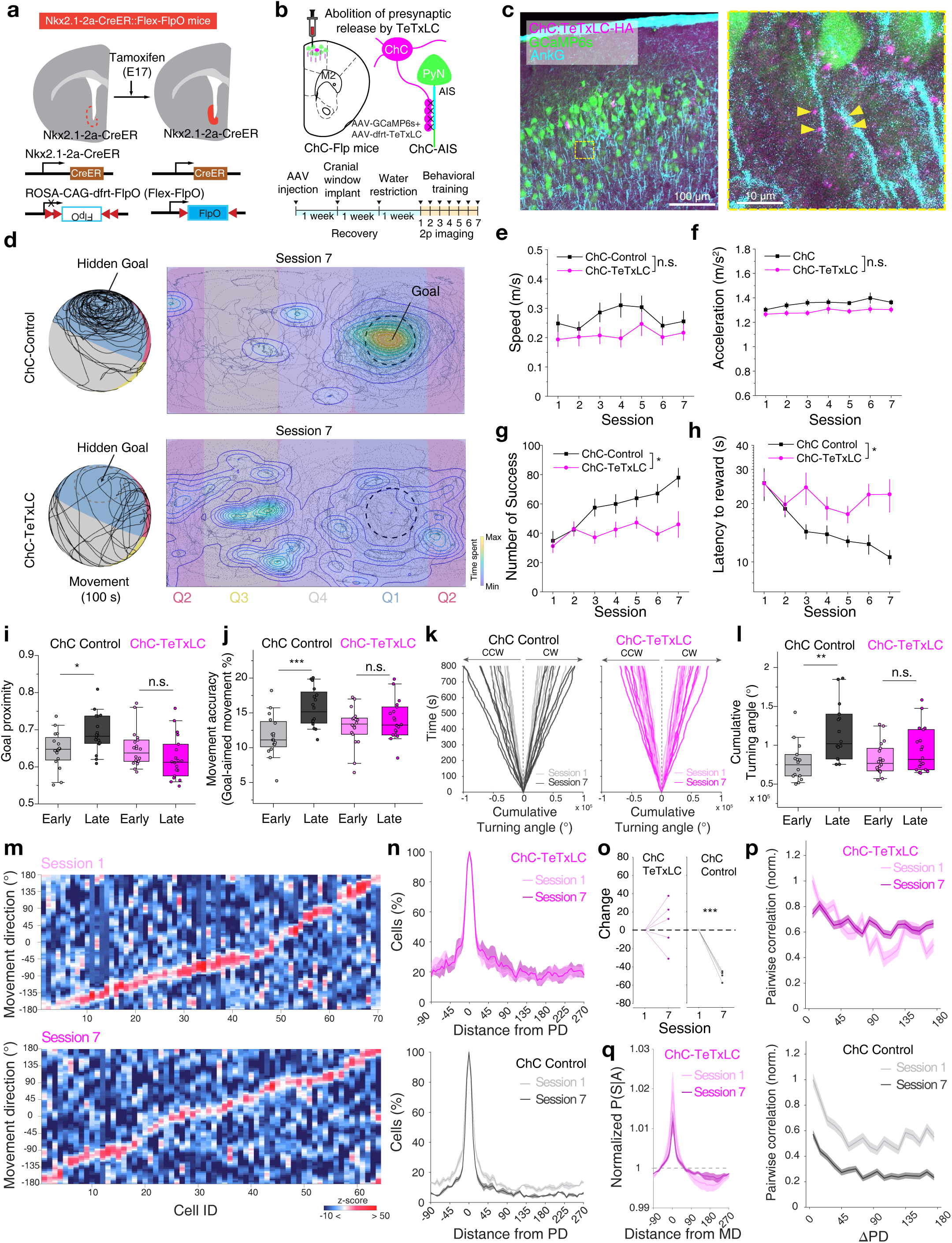
Abolition of ChC inhibition impairs the refinement of direction coding and motor learning. **(a)** Generation of Nkx2.1-2a-CreER::Flex-FlpO mice. Nkx2.1-2a-CreER::Flex-FlpO mice were generated by crossing Nkx2.1-2a-CreER with ROSA-Flex-FlpO mouse lines. The targeting vector containing Rosa26 homology arms, a CAG promoter, and a FLEX-Flp cassette was constructed. Similar to Nkx2.1-2a-CreER::Ai14, tamoxifen was administered to timed pregnant SW females by oral gavage at E17. **(b)** Schematic for selective abolition of GABA release from ChCs in M2 by expressing tetanus toxin light chain (TeTxLC) in ChC-Flp mice (Nkx2.1-2a-CreER::Flex-FlpO mice). **(c)** A representative image of ChC expressing TeTxLC-HA and neighboring neurons expressing GCaMP6s in Nkx2.1-2a-CreER::Flex-FlpO mice (left) and post-hoc validation of ChCs’ axonal projection to the axon-initial segment of neighboring PyNs by Ankyrin G (AnKG) staining (right). **(d-l)** Behavioral impact of ChC manipulation. **(d)** Representative movement traces of mice in ChC-TeTxLC (top) and ChC control (bottom) navigating on the multi-textured floating ball maze in Session 7 (100 s). A circle with a dashed line indicates the goal spot on the ball. Heat map and contour lines of the times of mice spent on the location are presented. **(e)** Average movement speed (n = 9 for ChC-TeTxLC group; n = 8 for ChC control group; two-way repeated measures ANOVA, *F_group_* = 1.16, *P* = 0.32). **(f)** Average movement acceleration (two-way repeated measures ANOVA, *F_group_* = 3.22, *P* = 0.12). **(g)** Average number of successes obtained in training sessions (two-way repeated measures ANOVA, *F_group_* = 8.54, *P* = 0.011). **(h)** Average latency to reward (two-way repeated measures ANOVA, *F_group_* = 7.81, *P* = 0.0136). **(i)** Average goal proximity (two-tailed *t*-test, *t* = −2.52, *P* = 0.017 for ChC control; *t* = 1.38, *P* = 0.177 for ChC-TeTxLC). **(j)** Movement accuracy (two-tailed *t*-test, *t* = −3.80, *P* = 6.67 × 10^−4^ for ChC control; *t* = −1.05, *P* = 0.30 for ChC-TeTxLC). **(k)** Cumulative turning angle over time in Sessions 1 and 7. **(l)** Comparison of cumulative turning angle between ChC-TeTxLC and ChC control (two-tailed *t*-test, *t* = −3.21, *P* = 3.51 × 10^−3^ for ChC control; *t* = −1.39, *P* = 0.175 for ChC-TeTxLC). **(m)** Example tuning curves of individual premotor neurons for movement direction in sessions 1 (top) and 7 (bottom) in ChC-TeTxLC condition. Data is sorted from the location of peak likelihood probability P(active|MD). **(n)** Normalized percentage of active cells in the population as a function of distance from the preferred direction (n = 378 cells for Session 1, n = 416 cells for Session 7 in ChC-TeTxLC; n = 665 cells for Session 1, n = 689 cells for Session 7 in ChC control). **(o)** Changes of the percentage of active cells from Session 1 to Session 7 (n = 5 mice for ChC-TeTxLC, two-tailed Student’s *t*-test. *t* = 0.56, *P* = 0.606; n = 4 mice for ChC control. *t* = −17.97, *P* = 3.76 × 10^−4^). **(p)** Pairwise correlations with respect to ΔPD normalized by Session 1 (angular difference in preferred direction between neuronal pairs, n = 15028 pairs for Session 1, n = 22589 pairs for Session 7 for ChC-TeTxLC; n = 57956 pairs for Session 1, n = 62104 pairs for Session 7 for ChC control). **(q)** Changes in posterior probabilities normalized by chance level as a function of distance from MD with learning. n.s., not significant; \**P* < 0.05; \*\**P* < 0.01; \*\*\**P* < 0.001; Error bars and shading indicate s.e.m. In the box plot, the midline, box size, and whisker indicate median, 25-75th percentile, and 10-90th percentile, respectively.

To determine the specific role of ChCs in suppressing irrelevant activity, we analyzed the probabilistic distributions of neural tuning to MD as a function of learning. Compared to the decrease in activity responding to non-PDs seen with learning, ChC-TeTxLC mice did not show a reduced likelihood of non-PD activity over training (**Fig. 7m**). While the estimated probability of the cell being active for non-PD movement (distance from PD > 90 degree) decreased in the ChC control mice with learning, such reduction was not observed in ChC-TeTxLC mice (**Fig. 7n**). When the MD was opposite to the cell’s PD, the probability of the cell being active significantly decreased in the ChC-control group, but not in the ChC-TeTxLC group (**Fig. 7o**). Consistently, we found that the decreases in pairwise correlation of neural responses with learning were specific to ChC control but not ChC-TeTxLC mice (**Fig. 7p**). Additionally, posterior probability density functions of MD were flattened over sessions in the ChC-TeTxLC group, indicating impaired encoding of movement direction due to the abolition of ChC-mediated inhibition (**Fig. 7q**). Given that intact ChC inhibition mediated the suppression of incorrectly tuned responses with learning, these results collectively suggest a necessary role of ChCs in refining distributed neural codes for active locomotion control.

### Heterogeneous axo-axonic synaptic plasticity underlies organized motor control

Our imaging data raised the possibility that ChC inhibitory synaptic strength may not be uniformly altered by learning. To understand the cellular evidence in support of this claim, we examined the structural rearrangement of axo-axonic synapses as proxies for the synaptic strength changes in target neurons. We developed an automated detection algorithm to discern the individual structures of ChC innervation on AISs of target PyNs, by using volumetric images of M2 (**Extended Data Fig. 7**, See Methods). A large number of inhibitory presynaptic and postsynaptic ChC-AIS structures, ChC axonal boutons, and gephyrin signals (GABAergic inhibitory synapse scaffolding protein) along the AISs were systematically analyzed in both the experimental group (n = 1634 AISs in 5 mice) and control group with *ad libitum* water (n = 3116 AISs in 6 mice) (**Fig. 8a-d**). We estimated structural synaptic strength by measuring presynaptic structural efficacy (Pre-SSE) as a presynaptic feature and postsynaptic structural efficacy (Post-SSE) as a postsynaptic feature based on ChC-AIS contacts and the corresponding gephyrin intensity, respectively (**Extended Data Fig. 8a**). The Pre- and Post-SSE distributions showed intrinsic heterogeneity in both control and experimental mice, indicating uneven inhibitory weights for connections between ChCs and M2 neurons (**Fig. 8d-h**). These imaging results are consistent with early Golgi staining and recent electron microscopy data that show non-random distribution of ChC axonal terminations ^18, 38^. The strong correlation between Pre- and Post-SSE showed the balance of presynaptic and postsynaptic strengths on AIS (**Fig. 8i**).

**Figure 8.**
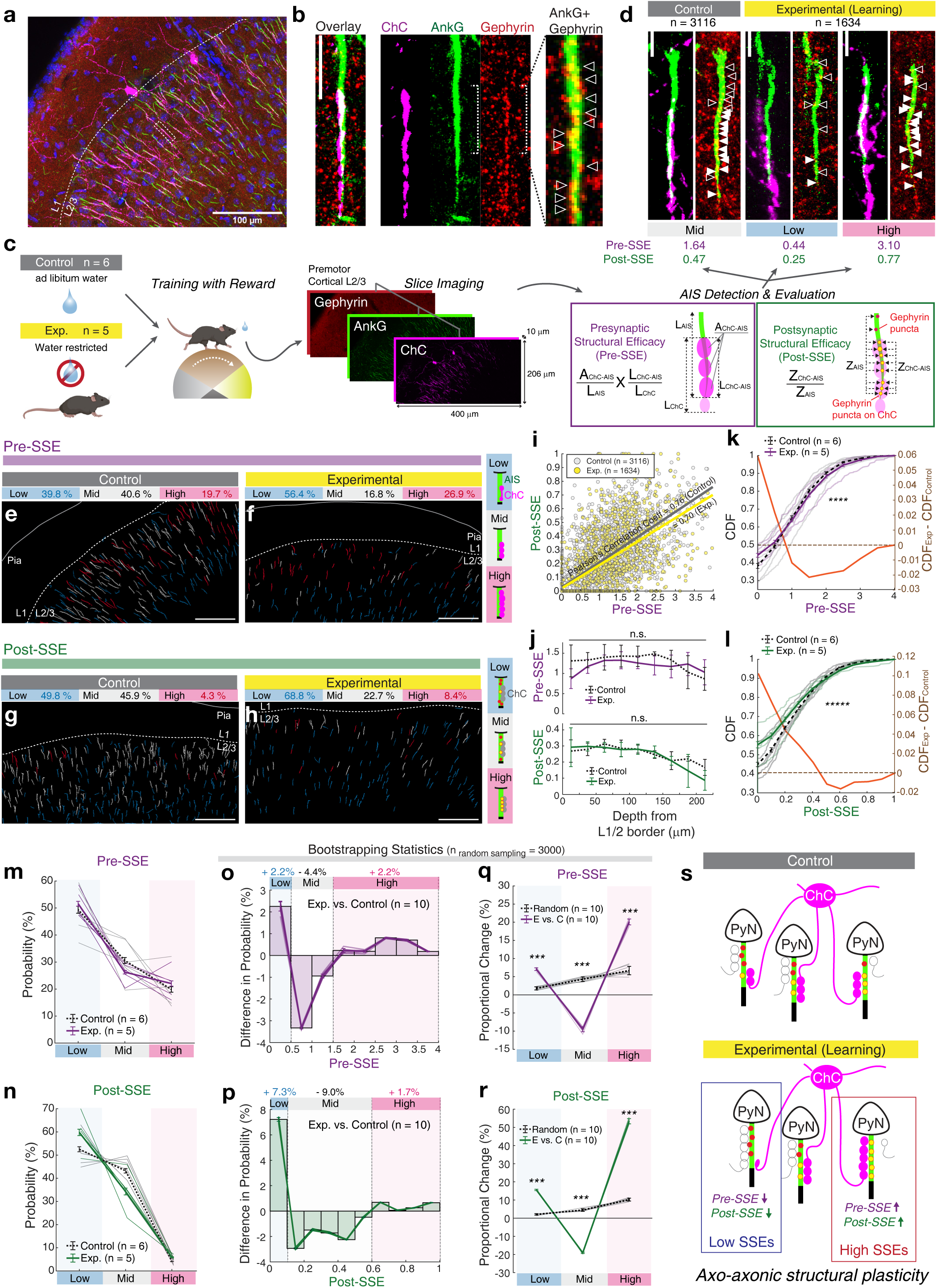
Heterogeneous axo-axonic synaptic plasticity underlies organized motor control. **(a, b)** Immunostaining of the axon initial segment (AIS) by AnkG (green), and inhibitory postsynaptic gephyrin (red) visualizes the corresponding synaptic composition (yellow) of ChC (magenta) to AIS contacts. DAPI stains nuclei (blue). **(c)** Schematics of automated detection of ChC-innervated AISs (ChC-AISs). **(d)** Representative examples of ChC-AIS evaluation of presynaptic and postsynaptic structural efficacy (Pre-SSE and Post-SSE). Filled and empty arrows indicate ChC-AIS and non-ChC-AIS synapses respectively. SSE values were classified into three subgroups (high: > 1.5 for Pre-SSE, > 0.6 for Post-SSE; low: < 0.5 for Pre-SSE, < 0.1 for Post-SSE; mid between high and low). **(e-h)** Examples of Pre-SSE and Post-SSE distribution in each slice for the experimental (learning) group and control group. **(i)** Scatter plot of Pre-SSEs and Post-SSEs for each group. **(j)** SSEs by the position of AISs within the cortical layer 2 for each group, which is measured by the depth from the border of layers 1 and 2. **(k, l)** CDFs from individual mice (*P* = 5 × 10^−5^ for **k** and *P* = 2 × 10^−11^ for **l**, error bars = s.e.m., left axis) and the difference between the averages for the experimental and control condition (Exp. vs. control, right axis). **(m, n)** Probabilities by Pre-SSE and Post-SSE strength subgroups (high, mid, and low) from individual mice. Error bars indicate s.e.m. **(o, p)** Differences of probability distributions for Pre-SSE and Post-SSE (Exp. vs. control) and **(q, r)** proportional changes (Exp. vs. Control) in the probability of each subgroup which is confirmed by 10 independent robust random samplings (n = 3000 each). In both SSEs, high and low subgroups were increased. The results were compared to proportional changes between unlabeled random pairs. For bootstrapping statistics (**o-r**), the mean and error of each bin were calculated from ten distributions generated by independent robust random sampling (n = 3000). For each random sampling, the probability distribution of a mouse was randomly selected from each condition (Exp. and Control) and used to calculate differences. Error bars indicate S.D., *P* = 2.0 × 10^−4^ for all bins. **(s)** Model for axo-axonic structural plasticity by heterosynaptic competition. The CDFs were tested by two-sample Kolmogorov-Smirnov test and the other distributions were tested by the two-tailed Wilcoxon-Mann-Whitney test. Scale bar = 100 μm (**a, e-h**), 5 μm (**b, d**). Every fluorescent image is presented by maximum intensity projection of the corresponding volumetric stack with pseudo-colors. \*\*\**P* < 0.001; \*\*\*\**P* < 0.0001; \*\*\*\*\**P* < 0.00001.

A subset of ChCs in the prelimbic cortex was reported to selectively control PyNs located more superficially (< 50 μm from L1/2 border) for their specific projection to the basolateral amygdala ^31^. We tested whether the heterogeneity among axo-axonic synapses depends on the cortical depth of AIS location. The overall synaptic strengths were not different by the location of AISs in L2/3 of control mice and experimental mice (0 – 250 μm from L1/2 border, dotted and solid lines in **Fig. 8j**). However, we found that high and low portions of SSE distribution increased in the experimental group, suggesting that the increased heterogeneity in ChC-PyN connections with learning is not solely based on the cortical depth of AIS location (**Fig. 8f, h**).

To decipher how learning reorganizes axo-axonic inhibition on AISs, the cumulative density functions (CDFs) of SSEs in the experimental group of mice were compared to the control group (**Fig. 8k, l**). If a subset of neurons is more tightly controlled by ChCs with learning (reflecting a sparse and determinant inhibitory control), we expect to see an increased percentage of AISs largely covered or scarcely covered by ChC boutons, respectively. Indeed, we observed an increased portion of both weak and strong ends in the distributions of pre and post-SSEs in the trained mice. Classifying AISs into three subgroups by their SSE strength (high, mid, and low), 4-9% of ChC-AISs underwent structural changes from mid to high or low (**Fig. 8m, n**). To validate whether this alteration was a result of training and not of high variance among samples (**Extended Data Fig. 8b-e**), we performed a bootstrapping test (n = 3000 random sampling) on the data from each mouse (**Extended Data Fig. 8f-i**). Obtained from multiple bootstrapping tests (n = 10), changes in the SSE distributions for trained mice showed increases in high (> 1.5 for Pre-SSE, > 0.6 for Post-SSE) and low (< 0.5 for Pre-SSE, < 0.1 for Post-SSE) value ranges (**Fig. 8o, p**). We found increased divergence only between the experimental and control groups (**Extended Data Fig. 9f-i**, Exp. vs. Ctrl.), and not between other condition comparisons (Exp. vs. Exp.; Control vs. Control; Unlabeled vs. Unlabeled). The alterations in each subgroup were analogous and tested as significant (**Fig. 8q, r**). Thus, these data suggest that ChCs differentially regulate the degree of inhibition applied to each axo-axonic synapse during learning (**Fig. 8s**).

## Discussion

Given the diversity of interneuron types and their specific connectivity in neural circuits, it has been a longstanding belief that the impact of inhibition on neural computation depends on a combination of interneuron cell-types and their connectivity patterns ^30, 39–41^. Genetically-defined interneuron groups such as PV, SOM, and VIP, and their connectivity motifs have profoundly advanced our understanding of cortical computation and psychiatric disorders. Different interneuron subtypes display their own circuit motifs, but subtypes remain widely heterogeneous. Therefore, we still do not fully understand the precise and unique roles of inhibition mediated by most single interneuron subtypes.

Despite the discovery of ChCs over four decades ago ^42^ and their clinical relevance to neuropsychiatric disorders such as schizophrenia characterized by GABAa receptor dysregulation at AIS ^43^, the functional role of ChCs in cognition and behavior has been enigmatic. Using a combination of chronic two-photon population imaging together with analyses on the structural plasticity at axo-axonic synapses, we demonstrated a critical role of the ChC in shaping the cortical coding of L2/3 M2 neurons during motor learning. ChC activity was necessary for improving the precision and accuracy of directional movement control. ChCs fine-tune the direction selectivity of individual premotor neurons and their collective tone. Notably, global population activity was not perturbed when ChCs were blocked. The necessary role of ChCs without perturbing overall network excitability implies that ChCs mediate their regulatory effect by establishing a specific neuronal connectivity pattern for learning. Quantification of a large number of ChC-AIS synapses uncovered revealed subsets of them underwent modifications associated with learning. This analysis revealed a significant degree of heterogeneity, which provides evidence of unequal inhibitory weighting among ChC-PyN connections during learning.

Diverse interneurons have been shown to be involved in motor learning by shaping neuronal population activity. Particularly, it has been shown that SOM-INs targeting the dendrites and spines of PyNs in the primary motor cortex undergo plasticity during motor learning and control learning-dependent sequential activation of PyNs in a manner comparable to what is attributed to ChCs here ^44^. In addition, inhibiting SOM-INs during new learning destabilized previous learning-dependent sequential activity of PyNs, suggesting the importance of SOM-IN inhibition for maintaining previously learned sequential activation patterns and behavioral improvement when a new motor skill task is introduced. However, in our study, we observed that acute optogenetic suppression of SOM-INs did not significantly disrupt navigation task performance after learning had been established (**Fig. 3a, Extended Data Fig. 4**). Given the reduced activity of SOM-IN activity during learning and partial restoration of naive-like activity by post-learning SOM-IN reactivation ^45^, the acute post-learning inhibition of SOM-INs is less likely to dominantly regulate neural representations formed with learning. We cannot rule out the possibility that SOM-INs are involved in shaping learning by dendritic-specific computation through synaptic plasticity on the dendritic spines of PyNs. It is possible that the suppression of irrelevant activity that is attributed to ChCs in the present study can be partly mediated by SOM-INs via their recurrent connectivity ^46^. Subpopulations of SOM-INs increase and decrease activities during motor learning in a task-specific manner ^44^. It is likely that diverse and task-specific activity of SOM-INs may mediate the temporal shift of motor learning-induced sequential activity in PyNs, which can complement organized population coding in sequence during goal-directed navigation.

The abolishment of perisomatic inhibition in PV-TeTxTC mice excessively increased the overall activities of M2 neurons, consistent with the notion that PV-INs are critical regulators of cortical excitatory and inhibitory (E/I) balance ^33, 34^. Perisomatic inhibition from PV-INs provides powerful inhibitory control over the PyN population ^29–32^. Fast-spiking PV-Basket cells (PV-BCs) are well suited for regulating the balance, gain and network oscillation of relatively broad PyN populations ^47, 48^. Balanced and delayed inhibition of fast-spiking INs with movement activity may provide temporal sharpening of motor command and suppress irrelevant activities ^35^.

PV-BCs and ChCs have different dendritic arborizations and laminar locations of their soma. They receive spatiotemporally distinct patterns of excitatory synaptic inputs from local PyNs, long-range thalamocortical connections, and neuromodulatory inputs, and in turn form different recurrent feedback excitation and inhibition patterns in a microcircuit ^49, 50^. The distinct wiring features and output properties of PV-BCs and ChCs may give their differential control over the spike-timing of PyNs ^51^. Different cholinergic modulatory effects of synaptic inhibition between ChCs and PV-BCs have been reported ^52^. Notably, a previous *in vivo* study showed that ChCs significantly increase their firing rate during arousal when brain states switched from slow to theta oscillations in the hippocampus and showed a low-amplitude desynchronized field potential in the prelimbic cortex, while the firing rates of PV-BCs and PyNs remained unchanged ^53^. Given that arousal is often associated with cholinergic signaling, differential modulation by cholinergic receptor activation between ChCs and PV-BCs may result in their distinct contributions to population activity control in a state-dependent manner. A future study is needed to further clarify the functional difference between ChCs and other types of BCs in behavior control, which is important to specify the role of perisomatic inhibitory circuit motifs.

Our findings support a model that ChCs exert differential strengths of control over subsets of PyNs after learning. There exists substantial variability in the magnitude of ChC input to PyN AISs ^54, 55^. This variability reflects the ability of ChCs to regulate their inhibitory strength based on the characteristics of its target cell, and may have a role in shaping the functional properties of PyNs. Variability in the number of axo-axonic ChC synapses correlated with structural features of individual target PyNs including laminar depth of the soma, other sources of perisomatic inhibition, and size of soma and AIS ^18^. The innervated PyNs are not distributed at random, but rather show a clustered distribution in a spatially heterogeneous fashion ^56^, supporting the existence of target selectivity and/or avoidance by ChCs on local PyNs. Two factors possibly select PyNs for stronger/weaker inhibition. First, ChCs differently inhibit PyNs based on the projection targets of the PyNs. Recent studies showed that a subset of L2 ChCs selectively innervates PyNs projecting to the basolateral amygdala over those projecting to the contralateral cortex in the prelimbic cortex ^31^. Second, ChCs differentially inhibit PyNs in an activity-dependent manner during learning ^57^. In this study, we analyzed a restricted subset of ChCs in the L2 and sampled neighboring PyNs in the shallow part of L2/3. Deeper L2/3 PyNs may have different projecting targets. Nevertheless, our findings on learning-dependent changes of axo-axonic synapses in the shallow L2/3 suggest that the difference in ChC-mediated inhibition is not solely resulted from the anatomical properties of target neurons. Given experimental evidence for the activity-dependent structural plasticity of axo-axonic synapses ^58–60^, it is conceivable that the plasticity of axo-axonic synapses provides an activity-dependent regulatory mechanism in the cell-by-cell level to fine-tune neuronal excitability across diverse cells involved in different functional networks. This can be mediated through precise inhibitory synaptic plasticity, which has been theoretically suggested as essential for the formation and maintenance of functional cortical circuitry ^61, 62^. Our findings suggest that the learning-dependent synaptic plasticity of axo-axonic synapses provides individualized inhibition to each PyN, which is adjusted to precisely balance excitation and inhibition based on the relevancy of its activity to the task goal.

Our measures of direction selectivity and population vector coding of MD may not fully capture high-order representations such as state-dependent action selection. It is presumed that multiple other inputs, such as cholinergic, thalamic, and other cortico-cortical inputs, innervate ChCs and PyNs. Premotor mechanisms of direction selectivity coding, movement direction, and inhibitory synaptic plasticity coming from different inputs remain to be studied.

Here we revealed a specialized inhibitory role for ChCs in experience-dependent plasticity of cortical coding, which allows for flexible and robust learned behavior. Given the unknown functions of a number of heterogeneous interneuron types, our results suggest that such heterogeneity may have evolved to serve specific functional needs. Unraveling the multitude of inhibitory effects on neural representation and signal transmission across all cell types is essential for understanding how cognitive functions are internally and externally governed to give rise to meaningful behavior. Precise balance and target-specific inhibitory synaptic plasticity by each inhibitory cell type are thought to contribute to the formation and retrieval of distributed memory, providing fundamental neural circuit architecture for learning. The heterogeneity of inhibitory cell types with their diverse receptors and innervation patterns offer us a glimpse into how the intricate and versatile neural coding in the cortex can be adopted to generate complex and flexible behavior. Our work sheds light on the importance of future studies to investigate additional single-cell type specific plasticity rules and their impacts on network activity and behavior.

## Methods

### Animals

C57BL/6J, PV-Cre (Cat.# 8069), SOM-Cre (Cat.# 13044) mice, Vipr2-Cre (Cat.#: 31332), Ai14 mice (Cat.# 7914) from Jackson laboratory (Bar Harbor, ME, USA) and Swiss Webster (Cat.#24) mice from Charles River Laboratory (Boston, MA, USA) were used in this study (4-9 weeks old, both sexes). All mice were maintained on a 12 hr light/ 12 hr dark cycle. All experimental procedures were carried out in accordance with protocols approved by Johns Hopkins University Animal Care and Use Committee, the Max Planck Florida Institute for Neuroscience Institutional Animal Care and Use Committee, and National Institutes of Health guidelines. For genetically targeting ChCs, Nkx2.1-2a-CreER and Vipr2-Cre mice were used. Swiss Webster mice were used as the background strain for Nkx2.1-2a-CreER mice while C57BL/6J mice were used as the background strain for Vipr2-Cre mice. Nkx2.1-2a-CreER and ROSA-Flex-FlpO mouse lines were generated in the laboratory of Hiroki Taniguchi at the Max Planck Florida Institute for Neuroscience. For Nkx2.1-2A-CreER mice, a 2A-CreER cassette was inserted in the frame immediately after an open reading frame of an Nkx2.1 gene. The targeting vector containing 5’ and 3’ homology arms, a 2A-CreER cassette, an frt-Neo-frt cassette, and an HSV-TK gene was constructed. For FLEX-Flp mice, the targeting vector containing Rosa26 homology arms, a CAG promoter, and a FLEX-Flp cassette was constructed. Both targeting vectors were generated using a PCR-based cloning strategy. 129SVj/B6 F1 hybrid ES cells (V6.5) were electroporated with the targeting vectors and subjected to drug resistance tests. Neomycin-resistant ES clones for Nkx2.1-2A-CreER and FLEX-Flp were screened by mini-Southern blotting and PCR, respectively, for correct targeting. Positive ES clones were used for tetraploid complementation to obtain male heterozygous mice following standard procedures. Nkx2.1-2a-CreER::Ai14 and Nkx2.1-2a-CreER::Flex-FlpO mice were generated by crossing Nkx2.1-2a-CreER with Ai14 and ROSA-Flex-FlpO mouse lines, respectively. Both created ChC mice showed no visible behavioral phenotypes and were able to learn the behavior task used in this study.

### Tamoxifen (Tmx) administration

Tmx was administered to timed pregnant Swiss Webster females that were bred to Nkx2.1-2a-CreER::Flex-FlpO males by oral gavage at E17 to induce CreER activity in the offspring. To achieve a high density of ChCs, the Tmx dose was adjusted to 3 mg/30 g of body weight. Tmx solution was prepared at a working concentration of 20 mg/ml in corn oil (SIGMA), kept protected from light, and kept refrigerated for no longer than one month.

### Animal surgery and stereotactic viral injection

Surgeries were performed on 4–8 week-old mice. Mice were anesthetized with an intraperitoneal injection of an anesthetic cocktail containing ketamine (87.5 mg/kg) and xylazine (12.5 mg/kg) (Santa Cruz Animal Health). The animal’s scalp was shaved and any remaining hair was removed with a hair remover lotion (Nair, Church & Dwight Co, Inc, Princeton, NJ, USA). Ophthalmic ointment (Puralube Vet Ophthalmic Ointment) was applied to prevent eyes from drying. Next, the animal was placed in a stereotaxic device (Kopf, Model 900 Small Animal Stereotaxic Instrument). The surgical region was scrubbed by 10% betadine solution (Purdue product LP, Stamford, CT, USA) and cleaned with 70% alcohol solution. Body temperature (37–38 °C) was maintained by a thermostatically controlled heating pad (Harvard Apparatus, Holliston, MA, USA). A small incision was made on the scalp. A small craniotomy (∼0.5 mm in diameter) was made above the injection site (the right premotor cortex, A/P: +1.75 mm, M/L: +0.3 mm from the bregma, D/V: −0.25 mm from the brain surface). AAV1-hSyn-GCaMP6s-WPRE-SV40 (400 nl, Penn Vector Core) for C57BL/6J mice as the experimental group; a mixture of AAV1-CaMKII0.4-Cre-SV40 (Penn Vector core), AAV1-Syn-Flex-GCaMP6s-WPRE-SV40 (Penn Vector core), and AAV9-CAG-Flex-tetanus toxin light-chain (produced from ViGene Biosciences, MD) (1:1:1 ratio, total injection volume: 800 nl) for C57BL/6J mice as CaMKII-TeTxLC; AAV9-EF1a-DIO-eNpHR3.0-eYFP-WPRE-hGH (400 nl, Penn Vector Core) for PV-Cre as PV-eNpHR and SOM-Cre mice as SOM-eNpHR; a mixture of AAV1-hSyn-GCaMP6s-WPRE-SV40 (400 nl) and AAV1-CAG-Flex-tdTomato-WPRE-bGH (400 nl, Penn Vector core) for PV-Cre mice as PV control; a mixture of AAV1-hSyn-GCaMP6s-WPRE-SV40, AAV1-CAG-Flex-tdTomato-WPRE-bGH, and AAV9-CAG-Flex-tetanus toxin light-chain (ViGene Biosciences, MD) (1:1:1 ratio, total injection volume: 800 nl) for PV-Cre mice as PV-TeTxLC; AAV1-hSyn-GCaMP6s-WPRE-SV40 (400 nl) for Nkx2.1-2a-CreER::Ai14 as ChC control; a mixture of AAV1-hSyn-GCaMP6s-WPRE-SV40 (300 nl) and AAV9-CAG-dfrt-hM4Di-mCherry-WPRE (500 nl) for Nkx2.1-2a-CreER::Flex-FlpO mice as ChC-hM4Di; a mixture of AAV1-hSyn-GCaMP6s-WPRE-SV40 (300 nl) and AAV9-CAG-dfrt-TeTxLC-HA-WPRE (500 nl) for Nkx2.1-2a-CreER::Flex-FlpO mice as ChC-TeTxLC; a mixture of AAV1-hSyn-GCaMP6s-WPRE-SV40 and AAVDJ-hSyn-FLEX-TeTxLC-P2A-dTom (1:2,000 dilution, a gift from Sandeep Robert Datta Lab)(1:1 ratio, total injection volume: 800 nl) for PV-Cre as sparse PV-TeTxLC; a mixture of AAV1-hSyn-GCaMP6s-WPRE-SV40 and AAVDJ-hSyn-FLEX-TeTxLC-P2A-dTom (1:1 ratio, total injection volume: 800 nl) for Vipr2-Cre mice as ChC-TeTxLC; AAV1-Syn-Flex-GCaMP6s-WPRE-SV40 (400nl, Penn Vector core) for Vipr2-Cre mice as ChC-GCaMP6; AAV1-CAG-Flex-TdTomato (400nl, Addgene, Cat.#: 28306) for Vipr2-Cre mice were used for virus injection. The viral constructs were injected via a beveled glass micropipette (tip size 10–20 μm diameter, Braubrand) backfilled with mineral oil. Flow rate (100-150 nl/min) was regulated by a syringe pump (World Precision Instruments). Following virus injection, skin adhesive (3M vetbond) was applied to close the incision site. General analgesia (Buprenorphine SR, 0.6 mg/kg) was injected subcutaneously and mice were monitored until they recovered from anesthesia. Around one week later, mice were anesthetized and hair was removed. The scalp was removed in a circular shape, and the surface of the skull was cleaned. A craniotomy (3.2 mm in diameter) was made and a glass cranial window (3 mm diameter, Warner Instruments, Hamden, CT, USA) was implanted on the virus injection site and a custom-made headplate was attached to the exposed skull with dental adhesive (C&B-Metabond, Parkell inc, Edgewood, NY, USA).

### Virus generation

To create AAV9-CAG-dfrt-hM4Di-mCherry-WPRE and AAV9-CAG-dfrt-TeTxLC-HA-WPRE, pAAV9-CAG-dfrt-hM4Di-mCherry-WPRE and pAAV9-CAG-dfrt-TeTxLC-HA-WPRE vectors were created by infusion of dfrt-hM4Di-mCherry and dfrt-TeTxLC-HA ‘R-products into pAAV9-CAG vectors. 293FT cells were transfected with pAAV-transfer vectors, PAD-Helper vectors and pSerotypeSpecific (AAV9) plasmids using a standard calcium phosphate method. After 70 hrs, cells were harvested in PBS and resuspended in 150 mM NaCl and 50 mM Tris, pH 8.5. After 3 freeze-thaw cycles, the virus solution was spun down at 4000 x g for 30 min and the supernatant was incubated in benzonase (100 U/ml) at 37 °C for 1 h. Another centrifugation step of 4000 x g for 25 minutes was followed by filtration of the supernatant through 0.45 μm and 0.22 μm. Next, the virus solution is put on top of a sterile Iodixanol gradient in an ultracentrifugation tube (15%, 25%, 40%, and 60% Iodixanol in 1 M MgCl_2_, 2.5 M KCl, 5 M NaCl in PBS). After centrifugation at 31,000 RPM for 6 hrs, the 40% Iodixanol band was harvested and stored in DPBS and concentrated using 100,000 concentrator vials (Amicon). The virus solution was aliquoted and stored at −80 °C until use.

### Behavior training in a floating ball maze

More than 2 weeks (19∼25 d) after virus injection, mice were trained to perform a tactile-based spatial navigation task while head-restrained on a custom-made spherical floating ball maze ^22^. Mice in the water-restricted group were water-restricted at 1 ml per day for a week before training starts. Mice in the control group were given *ad libitum* water access and, in addition, 2 ml of 10% sucrose were given 30 minutes before training. The floating ball maze was made of an air-supported 8” diameter spherical styrofoam ball with quadrants that have different tactile surface textures (ex. plain, grooved, striped, and grid). A simple auditory tone (8 kHz for 2 s) was delivered at the beginning and end of training. During navigation, mouse movement was monitored by a CMOS camera (Thorlabs) and ball movement was continuously monitored and recorded by a Bluetooth motion sensor (LPMS-B, LP research) placed at the center of the ball. Mice were trained to navigate to find a hidden goal location (surface area of 30 ° solid angle) in which a 10% sucrose reward (∼10 μl per a reward) was delivered if they were on the location. To facilitate animals’ exploration, a brief air puff (60 psi, 200 ms) was applied to their hind limbs if they did not move for 30 seconds. The reward was delivered via a lick port positioned in front of the mouse’s mouth and its timing was controlled by a solenoid valve (NResearch, West Caldwell, NJ, USA). To prevent excessive stay within the goal location and excessive access to the goal, a 3-second refractory period was assigned for the reward; Either a stay in the goal location or a re-visit to the goal location within 3 seconds from the last reward did not provide an additional reward. The behavioral setup was controlled by a custom-written code in MATLAB (Mathworks, Natick, MA, USA). Daily training sessions lasted 800 seconds and occurred for 7 days.

### Optogenetic inhibition of parvalbumin (PV) and somatostatin (SOM)-positive interneurons

We bilaterally injected AAV9-EF1a-DIO-eNpHR3.0-eYFP-WPRE-hGH (400 nl, Penn Vector Core) into the L2/3 M2 of PV-Cre mice or SOM-Cre mice and implanted fiber-optic cannulae (200 μm core, 0.37 NA, BFL37-2000, Thorlabs) with head plates. After two weeks of recovery from surgery and one week of water restriction, mice were trained on a floating ball maze for a session (800 s) per day for 7 days. After 7 days of training, mice were trained for a session and simultaneously received bilateral optogenetic inhibition using 589nm-light administration (continuous 800 s during the session, ∼10 mW power at the optic-fiber tip, MBL-FN-589, Changchun New Industries Optoelectronic Technology) through the fiber-optic cannulae. On the following day, mice were trained without optogenetic inhibition as a control.

### Chemogenetic inhibition of ChCs

We bilaterally injected a mixture of AAV1-hSyn-GCaMP6s-WPRE-SV40 (300 nl) and AAV9-CAG-dfrt-hM4Di-mCherry-WPRE (500 nl) into the L2/3 M2 of ChC-Flp mice. After 7 days of training, to suppress the activity of ChCs expressing AAV9-CAG-dfrt-hM4Di-mCherry in a FlpO-dependent manner, mice were administered i.p. with clozapine N-oxide (CNO, Tocris, Cat.#: 4936) 45 mins before a behavioral session and underwent the session. On the following day, mice were administered i.p. with saline as control and underwent the session.

### *In vivo* two-photon imaging

Imaging was conducted with a two-photon microscope (Bruker) using two pulsed Ti:sapphire lasers tuned to a wavelength of 920 nm (MaiTai HP DeepSee, Newport Spectra-Physics) for GCaMP6 signals or a wavelength of 1,045 nm (HighQ-2, Newport Spectra-Physics) for tdTomato and mCherry signals. The calcium imaging dataset was motion-corrected using a custom-written MATLAB code based on a full-frame cross-correlation image alignment algorithm. Regions of interest (ROIs) were semi-automatically drawn using a custom algorithm based on fluorescence intensity, cell size, and cell shape by visually inspecting movies, average of movies, and standard-deviation of movies, and then selecting neurons that showed fluorescence transients at least once during a session. The same area was imaged across training days to trace changes from same ensembles, but due to intrinsic tissue movement, neuronal activity was realigned when calcium signals were analyzed. All pixels within each ROI were averaged to create a fluorescence time series, *F.* Surrounding ‘neuropil ROIs’ were drawn from 0.3 μm outside of the border of each neuronal ROI to 10.0 μm outside. Neuropil ROIs excluded adjacent neuronal ROIs. Pixels within neuropil ROIs that showed apparent calcium transients exceeding 3 standard deviations of the difference in fluorescence between each neuropil ROI pixel time series and the neuronal ROI time series were excluded from further analysis. The remaining neuropil ROI pixels were considered as background fluorescence and averaged to create a time-varying background fluorescence trace. The time-varying baseline (*F0*) of a fluorescence trace was estimated by the following procedure: A preliminary baseline fluorescence time series, *Pre F0*, was loess smoothed with 120 frames. Preliminary *ΔF*, *Pre ΔF*, was obtained by subtracting *Pre F0* from *F*. Noise of Pre *ΔF* was estimated by subtracting loess smoothed *Pre ΔF* from the standard deviation of *Pre ΔF*. The offset of *ΔF* was determined by the mean of the distribution of *Pre ΔF* that did not exceed two times the noise. The baseline fluorescence trace, *F0*, was estimated by adding the offset to *Pre F0*. *ΔF/F* for neuronal ROIs was obtained by subtracting *F0* from *F* and dividing it by *F0*. The background fluorescence trace *ΔF/F* for neuropil ROIs was estimated in the same manner, i.e., as the *ΔF/F* for neuronal ROIs. The background fluorescence trace was subtracted with a 0.7 weight from the neuronal ROI fluorescence trace for each neuronal ROI. For pixel-based imaging analysis of MD, fluorescence change maps of images were color-coded by MD and averaged. For pixel-based analysis of different movement paths, fluorescence change maps of images for each movement path were differently color-coded and overlaid.

### Tissue fixation, immunohistochemistry, and acquisition of confocal and light-sheet microscope images

Animals were deeply anesthetized with a mixture of ketamine and xylazine and then perfused transcardially, first with phosphate buffered saline (PBS, pH 7.4) and next with 4% paraformaldehyde (PFA) dissolved in PBS. The brains were removed and postfixed in 4% PFA overnight at 4 °C. For confocal microscope images, the brains were embedded in 10% melted gelatin solution for 50 minutes at 50°C. Melted gelatin solution was then refreshed and the gelatin solution with the embedded brains was solidified at 4 °C for about 30 minutes. Then, the solidified gel was trimmed around the embedded brain to a small cube shape and the cube was stored into 4% PFA overnight. The gelatin cube brain was coronally sectioned (100 μm thick) using a vibratome (Leica Biosystems, Buffalo Grove, IL, USA). For parvalbumin immunohistochemistry, after blocking with a solution (1 % Normal Donkey Serum/0.1 Triton-X in PBS), brain slices were incubated in primary antibodies (1:500 mouse anti-parvalbumin, #P3088, Sigma) for 48 hours at 4 °C. After washing 3 times in PBS, the brain slices are then incubated in a secondary antibody (Cy5-conjugated donkey anti-mouse IgG, Cat.#: 715-585-150, diluted 1:1,000 in PBS, Jackson ImmunoResearch Laboratories) for 2 hours at room temperature, followed by 3 washes in PBS. The brain slices were mounted on the slide glass with a DAPI-containing mounting solution (DAPI Fluoromount-G, SouthernBiotech, USA). For Nkx2.1-2a-CreER::Ai14 and Nkx2.1-2a-CreER::Flex-FlpO mice immunohistochemistry, after intraperitoneal administration of Ketamine/Xylazine mixture (50 mg/kg Ketamine, VETCO, 5 mg/kg Xylazine, AKORN) and a foot pinch to check adequate sedation, the mice were trans-cardially perfused with 15 ml cold saline solution followed by 20 ml of cold 4% PFA solution in PBS. After post-fixation in 2% PFA in PBS overnight, 60 μm brain sections were prepared using a Leica vibratome and collected in PBS, permeabilized in 0.5% TritonX-100 in PBS followed by washing for 1 h in blocking reagent (BR: 0.1% TritonX-100, 10% donkey serum in PBS). The sections were treated in primary antibody solution in BR overnight at 4°C. The following primary antibodies were used: rat anti HA (1:500, Roche, #11-867-423-001), chicken anti GFP (1:800, ABCAM, Cat.#: ab13970), and mouse anti AnkyrinG (1:500, UC-Davis/NIH NEUROMAB, Cat.#: clone N106/36 75-146). After three washes (10 min) in PBS, the slices were incubated in BR with secondary antibodies (Jackson ImmunoResearch) for 2 h at room temperature containing appropriate fluorophores (CyX or Alexa) at 1:1000. The slices were embedded in FluoromountG (Thermo Fisher) [K1] after two final washing steps in PBS. The fixed and immuno-stained brain slices were imaged using an upright confocal microscope (Zeiss LSM880, Germany). For light-sheet microscope images, brains were perfused as described above and were coronally sectioned (2 mm thick) using a vibratome (Leica Biosystems, Buffalo Grove, IL, USA). The brain slices were cleared using the Passive Clarity protocol (Yang et al., 2014). Images of the cortex were collected with a Zeiss Light-sheet microscope at 25x magnification. Light-sheet microscope image processing and 3D rendering were performed with arivis Vision4D.

### Preparation for acute cortical slices

Mice were anesthetized with isoflurane and decapitated. The brain was removed and rapidly placed into ice-cold cutting solution containing (in mM): 215 Sucrose, 26 NaHCO3, 20 Glucose, 4 MgCl2, 4 MgSO4, 2.5 KCl, 1 CaCl2, and 1.6 NaH2PO4. Cortical slices (300 μm thick) were prepared using a VT1000S vibratome (Leica). Slices were incubated at 32 °C for 30 minutes in a holding chamber filled with artificial cerebrospinal fluid (ACSF) containing (in mM): 124 NaCl, 26 NaHCO3, 10 glucose, 3 KCl, 1.25 NaH2PO4, 2.5 CaCl2, and 1.3 MgSO4. Slices were then placed at room temperature and allowed to recover for 1 hour before recording. All solutions were equilibrated with carbogen (95% O2/5% CO2).

### Electrophysiology

Whole-cell patch-clamp recordings (electrode resistance 5-9 MΩ) were performed at room temperature using a Multiclamp 700B amplifier (Molecular Devices). Layer 2/3 chandelier cells were patched in current-clamp configuration using potassium-based internal solution (in mM: 10 NaCl, 1 MgCl2, 4 NA2-ATP, 124 K-gluconate, 10 Na2-phosphocreatine, 16 KCl, 0.4 Na-GTP, 10 HEPES) in ACSF containing (in mM: 124 NaCl, 26 NaHCO3, 10 glucose, 3 KCl, 1.25 NaH2PO4, 2.5 CaCl2, and 1.3 MgSO4). To examine the effect of DREADD virus hM4Di, we recorded the firing properties of hM4Di-expressing neurons before and after applying 10 μM clozapine N-oxide (CNO, Tocris, Cat.#: 4936) (in ACSF).

### Data acquisition and analysis

All data analysis was performed using Excel (Microsoft), and custom codes and toolboxes in MATLAB. Behavioral sessions in which the mouse made more than three successful accesses to the goal were analyzed. The movement signals of the ball maze including quaternion and angular velocity were sampled at 100 Hz and smoothed using a lowess regression of 500 ms width. We extracted animals’ virtual coordinates on the ball relative to the center of the hidden goal spot. We acquired mouse movement-related information based on translational movement (forward and backward; right and left) by using a precise Bluetooth motion sensor (LPMS-B, LP research, Tokyo, Japan).

### Identification of movement bouts

We classified each behavior frame (sampled at 100 Hz, 1 ms bin size) as occurring during either rest or movement. We defined rest frames as those in which the movement speed is less than 0.02 m/sec and acceleration is less than 0.2 m/sec. We defined movement frames as those in which a movement speed is greater than 0.03 m/sec and not classified as rest frames. Parameter values were determined by visually comparing videos with movement speed and acceleration traces. The uncategorized frames which do not meet the above criteria were further categorized as following: If both ends of the bout of uncategorized frames were connected to rest frames and the duration of the bout of uncategorized frames was less than 1 second, the uncategorized frames were identified as rest frames. If either both ends of the bout of uncategorized frames were connected to movement frames or the state of connected frames of one end of the bout was different from that of the other end, the uncategorized frames were identified as movement frames.

### Movement direction

Movement direction was determined by the inverse tangent of forward-backward speed over right-left speed. Goal-angle was defined as the angle passing through the goal spot from the mouse’s two-dimensional virtual position. Goal-directed movement epochs were determined if the mouse’s movement direction was within the goal-angle. Percentage of goal-directed movement was determined by the ratio of goal-directed movement epochs to overall movement epochs. Goal-directed movement bout was identified as a bout of movement in which mice pursued the hidden goal location and received the reward. Non-goal-directed movement bouts were defined as bouts wherein the animal does not reach the goal.

### Classification of movement-related neurons

Modulation of neural activity was frequently observed between movement and rest epochs. We classified individual neurons into three categories, movement-positive, movement-negative, and non-movement. Movement positive or negative neurons were identified if their mean activity during movement epochs was significantly higher or lower than during rest epochs, respectively. If there was no significant difference in mean neuronal activity between movement and rest epochs, neurons were classified as non-movement cells.

### Estimation of probabilistic neural tuning curves

To estimate calcium events, calcium signals were deconvolved using an Online Active Set method to Infer Spikes (OASIS) with FOOPSI approach and auto-regressive models with order 2. The values of calcium event estimate *s* were threshold to 0, producing binary events that represent estimated calcium events. To focus on the tuning curves of premotor neurons to movement direction, we therefore excluded periods of immobility. Using this event estimate, we computed the probability of a neuron to be active (*active*) as the period of activity of a neuron over the total period of a recording session, which corresponds to the marginal likelihood probability distribution in a Bayesian approach. We divided the movement direction in 5° bins. Each bin represents a discrete state of movement direction. We then computed the likelihood probability distribution that a cell is active given movement direction, (*active*|*MD_i_*), as the period of activity in direction state *MD_i_* over the total period in state *MD_i_*. The distribution was smoothed with a Savitzky-Golay filter. To test statistical deviation from random firing patterns, we carried out a bootstrapping procedure as follows. For each cell, deconvolved data were circularly shifted by different random values of shifts and the likelihood probability distribution of the randomized data was estimated. This procedure was repeated 100 times with different random values of shifts. The mean value and standard deviation of the randomized dataset were used to compute the z-scored likelihood distribution of actual data. A cell is defined as an active cell if the likelihood distribution of actual data is significantly deviated from that of the randomized dataset (*P* < 0.05).

Neurons were considered to be directionally tuned if they showed significant tuning with movement direction (ANOVA, *F* test, *P* < 0.05) and they showed a good fit of tuning function. To obtain the tuning curve, the z-scored likelihood distribution of actual data was fitted with the sum of two von Mises distributions ^63^:

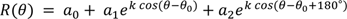

where R is the response to movement in a direction *θ*, *θ*_0_is the preferred direction, and a0, a1, a2, and k are fitting parameters. We estimated fitting parameters by using lsqcurvefit in the Matlab function. The preferred direction *PD_i_* for neuron *i* was defined as the movement direction corresponding to a peak in the tuning curve response. Neurons were considered to be direction-selective if they fit the von Mises distribution well (*P* < 0.05) and if their direction-selective index (DSI) defined as 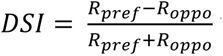 was greater than 0.4, where *R_pref_* and *R_oppo_* are the mean of the normalized response to the preferred direction and that of the opposite direction (the preferred direction + π), respectively.

### Bayesian decoding analysis

In addition to the estimation of the likelihood distribution, (*active*|*MD_i_*), we computed the probability of being in a given state of movement direction *MD_i_*, (*MD_i_*), as the period in the direction *MD_i_* over the total period of a recording session, which corresponds to the prior probability distribution. Using Bayesian approach ^64^, we inferred the posterior probability distribution function that the animal is in a movement direction given neuronal activity at a time *t*.

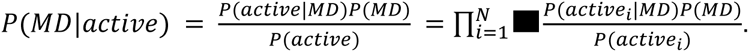

where *P*(*MD*|*active*) is a vector of a posterior probability distribution of behavioral states and *N* corresponds to the number of neurons used. To decode the direction of movement, we considered the direction associated with the maximum *a posteriori* (MAP).

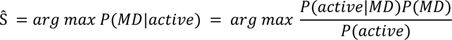

where Ŝ is the decoded movement direction. The decoding error was computed as the angular difference between the decoded and actual movement direction at the observed time.

### Pairwise correlations

We calculated pairwise correlations between neuronal pairs by measuring coincident events ^65^. For neuron *i*, the events *E_i_* were vectorized in a binary manner. The pairwise correlation for the pair *i* and *j* is given by

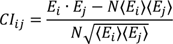

where *N* is the number of image frames, and 〈*E_i_*〉 is the expected number of events for neuron *i*. We categorized positively and negatively correlated pairs if the CI is greater than 0 and less than 0, respectively.

### Automated detection of ChC-AIS

To label ChCs in M2, we injected AAV1-CAG-Flex-TdTomato (400nl, Addgene, Cat.#: 28306) into L2 of M2 in Vipr2-Cre mice. Once the brain slices were stained with immunohistochemistry and imaged through the confocal microscope described in the tissue fixation, immunohistochemistry, and acquisition of confocal and light-sheet microscope images section, each stack of tiled confocal images (generally 3800 X 1950 X 10 in pixel or 400 X 206 X 10 in μm) from three different fluorescent channels (green for gephyrin puncta antibody, red for TdTomato expressed in ChC projection, and far-red for AnkG antibody targeting AIS) was processed simultaneously to detect AISs innervated by axonal cartridges of ChC and colocalized gephyrin puncta using a custom-written program in Matlab. Features of processed images were segmented automatically by boundary detection (bwboundaries function, Matlab) in each planar image of red and far-red channels after binarization by adaptive threshold (adaptthresh function, Matlab, sensitivity = 0.08 and neighborhood size = 19 for AIS, sensitivity = 0 and neighborhood size = 15 for ChC). Segments in different axial planes were then filtered through several criteria including intensity, size, and width and registered to recover the whole AISs or ChC cartridges. Only AISs longer than 10 µm were selected for the further analysis. For each registered AIS, we overlaid its segment on detected ChC cartridges in each axial plane and classified it as a ChC-AIS when any colocalized area existed. For demonstration purposes, fluorescence images shown in figures are maximum intensity projections of corresponding volumetric stack with pseudo-colors.

### Quantification of characteristic parameters

For each detected ChC-AIS segment, we calculated the lengthwise axis. Projected intensity profiles of ChC and AIS on the axis were then applied to measure the total AIS length (L_AIS_), the ChC cartridge length (L_ChC_), and the AIS length covered by ChC (L_ChC_AIS_). To ensure the exact L_ChC_ measurement, we extended the ROI 5 µm longer than the detected AIS segments we identified. Half of the maximum intensity for each profile was used as a threshold for measurement. The area of ChC on AIS (A_ChC-AIS_) was calculated as the mask size for segmentation. We computed the normalized ChC-AIS size, the presynaptic parameter, as

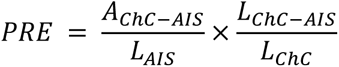

from these morphological quantities.

We used the particle detecting algorithm u-track ^66^ to identify gephyrin puncta in each plane of the green channel. This algorithm fitted Gaussian kernels approximating the two-dimensional point spread function of the microscope (σ = 1.5 pixel or 234 nm) around local intensity maxima, where both position and amplitude were free parameters in the fit. The intensity of detected gephyrin punctum is represented by a modified z-score calculated from the median and standard deviation of all the detected gephyrin puncta from the tiled image.

To evaluate the postsynaptic strength, a detected gephyrin punctum was classified as AIS-associated when its detected position resides in the boundary of a registered AIS. In the same manner, an AIS-associated gephyrin punctum was labeled ChC-AIS associated when its position colocalizes in the boundary of ChC on AIS simultaneously. Summation of the z-score of gephyrin puncta associated with the total AIS (Z_AIS_) and only a portion of ChC-AIS (Z_ChC-AIS_) was used for calculation of the corresponding gephyrin intensity, the postsynaptic parameter, as follows.

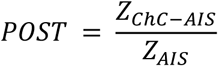

The corresponding gephyrin intensity equals 1 when all the gephyrin puncta on the AIS are located on the corresponding ChC. When the ChC does not have corresponding gephyrin puncta on the bound AIS, its value is 0.

## Statistics

Statistical analyses were performed using either MATLAB (Mathworks) or OriginPro (OriginLab). All statistical tests are described in the corresponding figure legends. Normality of distributions was tested for each dataset using Lilliefors test to decide whether to use parametric or non-parametric tests. For parametric tests, two-tail Student’s t-test, two-sample t-test, two-tailed paired t-test, one-way ANOVA, one-way repeated measures ANOVA, and two-way repeated measures ANOVA were performed to compare population means across animals and conditions except where noted. For ANOVAs, Fisher post hoc tests were used for multiple comparisons, When sphericity is violated, we used Greenhouse-Geisser correction. Chi-square analyses were used to compare proportions. Pearson’s linear correlation coefficient was used to measure the correlation between two variables. For non-parametric tests, two-tailed Wilcoxon-Mann-Whitney test and Friedman test with Dunn post hoc tests were performed to compare the population median and differences with dependent variables across animals and conditions, respectively. Two-sample Kolmogorov-Smirnov test was performed to compare probability distributions between groups. The number of animals and the number of cells used for analysis are specified in the figures, the figure legends, and the text.

## Data availability

Additional data that supports the findings of this study are available from the corresponding author upon reasonable request.

## Code availability

The code for semi-automated ROI selection can be found at https://github.com/fitzlab/CellMagicWand. Custom MATLAB scripts used in this study are available from the corresponding author upon reasonable request.

## Acknowledgments

We thank members of the Kwon laboratory for helpful discussions; R.Smith, N. Schappaugh, and K.S.Lee for comments on the manuscript. We thank D.J.Wendler and M.An for helping viral injection and brain fixation and K.S. Lee for helping brain clearing. GCaMP6s virus was available from the Genetically-Encoded Neuronal Indicator and Effector (GENIE) Project and the Janelia Farm Research Campus, specially V.Jayaraman, R.A.Kerr, D.S.Kim, L.L.Looger, and K.Svoboda. AAV-hSyn-FLEX-TeLC-P2A-dTomato was kindly provided from Sandeep Datta lab (Harvard Medical School). This work was supported by Max Planck Florida Institute for Neuroscience (to H-B.K., D.F., and H.T.), the National Institutes of Health Grants R01MH107460 (to H-B.K.), DP1MH119428 (to H-B.K) and R01MH115917 (to H.T.).

## Author contributions

K.J. and H-B.K. conceived and designed the study. K.J. designed and optimized head-fixed ball maze system. K.J performed most of the experiments including viral injection, electrophysiology, optogenetic manipulation, two-photon imaging and behavioral training. K.J. wrote behavior and calcium imaging data analysis programs and analyzed all data. M.C. and D.C performed immunostaining and data analysis. M.C. developed an algorithm detecting ChC-AIS synapses. B.B. helped with virus injection surgery. A.S. and H.T. made Flex-FlpO mice and provided Nkx2.1-2a-CreER::Ai14 and Nkx2.1-2a-CreER::Flex-FlpO mice. Y.O. and H.T. generated Nkx2.1-2a-CreER mice. D.F. and H.T. provided advice on the manuscript organization and data interpretation. K.J. and H-B.K. wrote the manuscript.

---

**Extended Data Figure 1.**
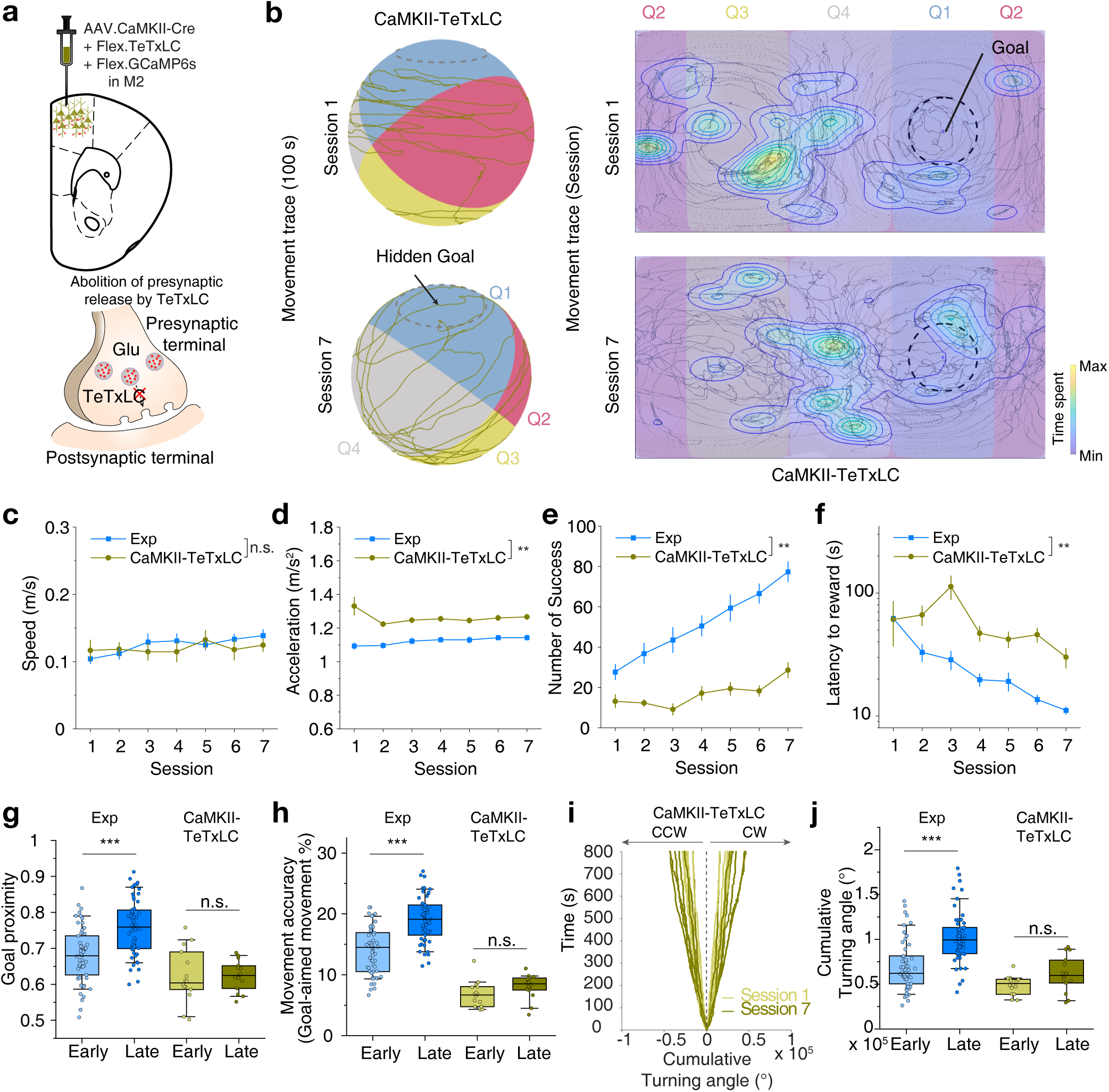
The premotor cortex is required for organized purposive motor control. **(a-j)** Blockage of local glutamate release in the premotor cortex impaired organized motor control. **(a)** Schematic of virus injection of tetanus toxin light-chain, which blocks local glutamate transmission from excitatory neurons in the premotor cortex by co-expressing AAV1.CaMKII-Cre and AAV1.Flex-tetanus toxin light-chain for CaMKII-TeTxLC group. **(b)** Representative movement trace (100 s) of a mouse in CaMKII-TeTxLC group on the ball (left) from training sessions 1 to 7 and its 2-dimensional projection (right). **(c)** Average movement speed of CaMKII-TeTxLC (n = 27 for the experimental group, Exp; n = 7 for CaMII-TeTxLC; two-way repeated measures ANOVA, *F_group_* = 0.45, *P* = 0.53). **(d)** Average movement acceleration (two-way repeated measures ANOVA, *F_group_* = 19.59, *P* = 0.0044). **(e)** Average number of successes obtained in training sessions (two-way repeated measures ANOVA, *F_group_* = 24.11, *P* = 0.0027). **(f)** Average latency to reward of CaMKII-TeTxLC (two-way repeated measures ANOVA, *F_group_* = 24.11, *P* = 3.50 × 10^−4^). **(g)** Average goal proximity of CaMKII-TeTxLC (two-tailed *t*-test, *t* = −5.19, *P* = 1.03 × 10^−6^ for Exp; *t* = −0.06, *P* = 0.95 for CaMKII-TeTxLC). **(h)** Movement accuracy of CaMKII-TeTxLC (two-tailed *t*-test, *t* = −7.04, *P* = 2.01 × 10^−10^ for Exp; *t* = −1.54, *P* = 0.135 for CaMKII-TeTxLC). **(i)** Cumulative turning angle of CaMKII-TeTxLC over time in Sessions 1 and 7. **(j)** Comparison of cumulative turning angle between Exp and CaMKII-TeTxLC (two-tailed *t*-test, *t* = −5.84, *P* = 5.76 × 10^−8^ for Exp; *t* = −2.05, *P* = 0.050 for CaMKII-TeTxLC). n.s., not significant; \*\**P* < 0.01; \*\*\**P* < 0.001; Error bars indicate s.e.m. In the box plot, the midline, box size, and whisker indicate median, 25-75th percentile, and 10-90th percentile, respectively.

**Extended Data Figure 2.**
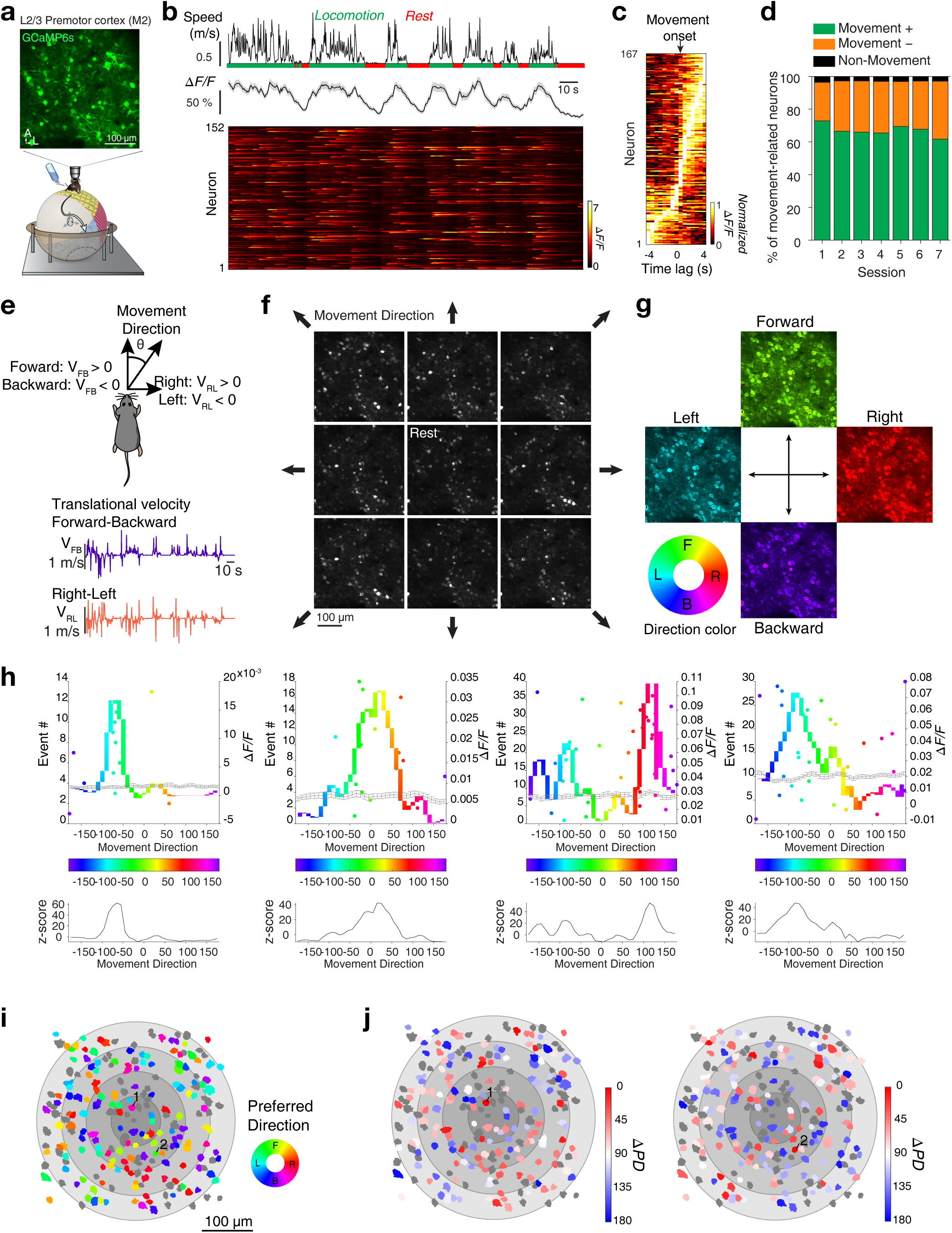
Movement direction selective responses of premotor neurons. **(a)** Schematic of the navigation task on ball maze with 2-photon calcium imaging and a representative field of view of layer 2/3 premotor neurons expressing GCaMP6. **(b)** An example trace of mouse movement speed (top), aligned averaged fluorescence transients (middle), and a corresponding heat-map raster plot (bottom). **(c)** Normalized fluorescence transients of premotor neurons aligned to movement onsets. **(d)** Summary graph of movement-related neurons throughout training. **(e)** Schematic that depicts the estimation of movement direction based on forward-backward and right-left speeds. **(f)** Average fluorescence imaging frames during responses to varying movement directions. **(g)** A color-coded, pixel-based map of neuron activity with respect to movement direction tuning. **(h)** Top, four example calcium transient events by movement direction (dots) and direction tuning curves of premotor neurons (top, colored-curve). Movement direction is color-coded. The gray line indicates the average tuning curve from shuffled data. Bottom, corresponding z-scores of the actual tuning curves normalized by the tuning curves of the shuffled data. **(i)** An exemplary preferred direction map. The color indicates the preferred direction of individual cells. Black and gray color indicate non-direction-tuned neurons. The salt-and-pepper layout of color indicates functional heterogeneity in direction-selectivity in the premotor cortex. **(j)** Two examples of the heterogeneous spatial distribution of ΔPD (angular difference in preferred direction) between neuronal pairs. The same neuronal population as in **i**, with color-coding by ΔPD (angular difference in preferred direction between reference neuron and another direction-tuned neuron). Numbered neurons indicate reference neurons.

**Extended Data Figure 3.**
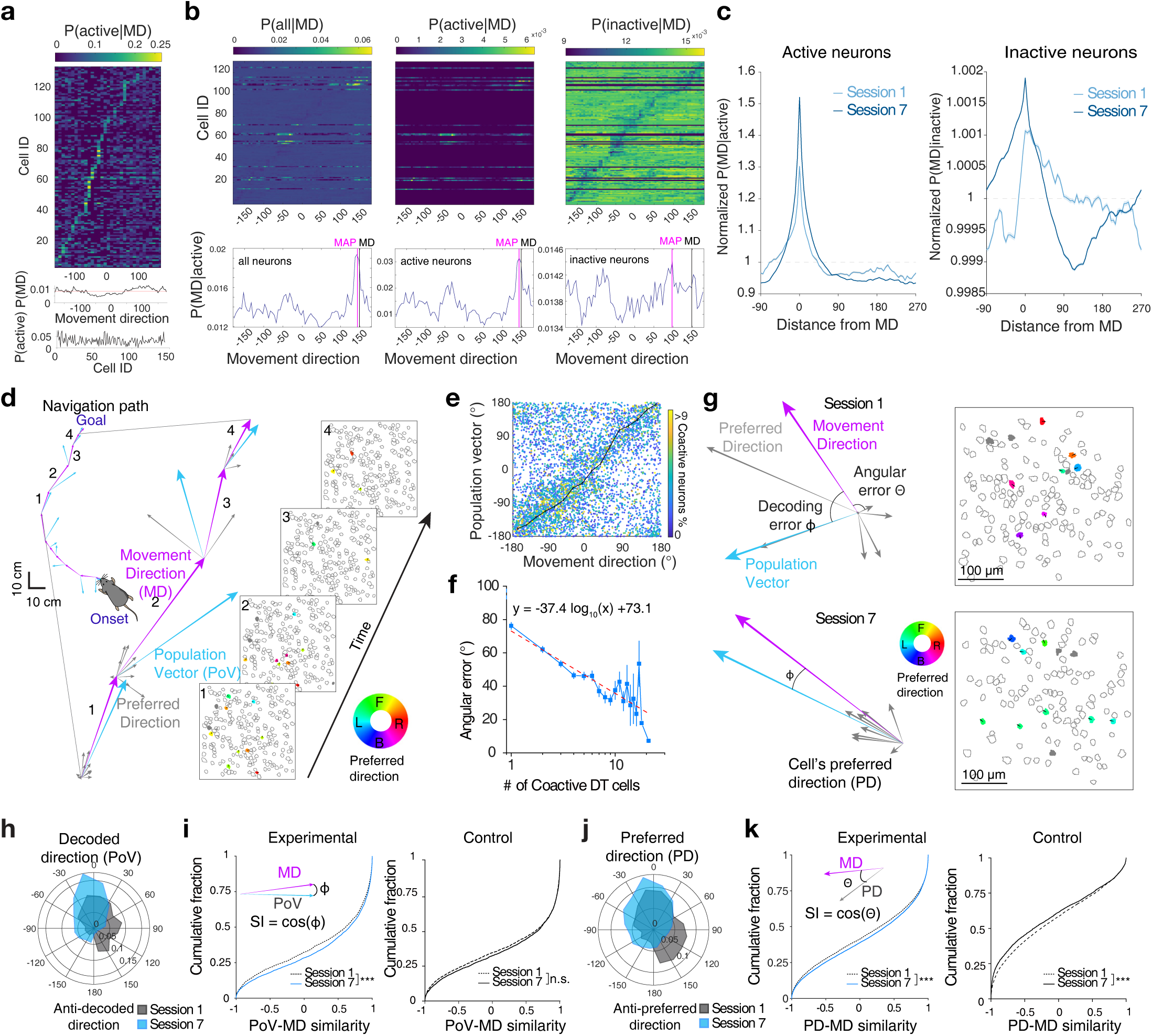
Population coding of movement direction in premotor neurons. **(a)** Movement direction tuning curves for each individual premotor neuron sorted from peak probability location (top). The prior probability of movement direction (middle). The marginal likelihood of being active (bottom). **(b)** Tuning curves of neurons corresponding to movement direction at a given moment (top) and posterior probability of movement direction (bottom) given activity from all neurons (left) from active neurons (center), and from inactive neurons (left). Actual movement direction (MD) and decoded movement direction estimated with maximum a posteriori (MAP) are shown. **(c)** An example of changes in posterior probabilities, *P(MD|A)*, normalized by chance level (dotted line) from a mouse of the experiential group, for active (left) and inactive neurons (right) as a function of distance from movement direction with learning. **(d)** Sparse population coding of movement direction during navigation; animal’s movement direction (MD, magenta), preferred directions of individual active direction-tuned neurons (PD, gray), and population vector (PoV, the vector sum of the preferred directions, blue) are superimposed. Corresponding maps of active neurons during movement. **(e)** Comparison between movement direction and sparse population vector. Each dot is color-coded by the percentage of coactive neurons in the population. **(f)** The angular error between the population vector and actual movement direction as a function of the number of coactive direction-tuned (DT) cells. The dotted line with a negative slope coefficient indicates a linear fit of the data. **(g)** An example of population coding of movement direction in session 1 (top) and session 7 (bottom). Magenta, cyan, and gray arrow lines indicate an animal’s movement direction, population vector direction, and active neurons’ preferred direction, respectively (left). Preferred direction map of the active population (right, filled ROIs). The filled color and ROI arrow indicate the corresponding preferred direction. **(h)** Polar distribution of angular errors between PoV and MD in the experimental group (session 1, gray; session 7, light blue). **(i)** Changes of the cumulative distribution of similarity index between PoV and MD with learning (Two-sample Kolmogorov-Smirnov test, D = 0.0495, *P* = 0.0003 for the experimental group, n = 6 mice; D = 0.027, *P* = 0.12 for the control group, n = 5 mice). **(j)** Polar distribution of angular errors between PD and MD in the experimental group (session 1, gray; session 7, light blue). **(k)** Changes of the cumulative distribution of similarity index between active neurons’ PD and MD with learning (Two-sample Kolmogorov-Smirnov test, D = 0.0533, *P* < 2.68×10^−10^ for the experimental group, n = 6 mice; D = 0.0584, *P* < 2.61×10^−10^ for the control group, n = 5 mice). ***P < 0.001; n.s., not significant. Error bars indicate s.e.m.

**Extended Data Figure 4.**
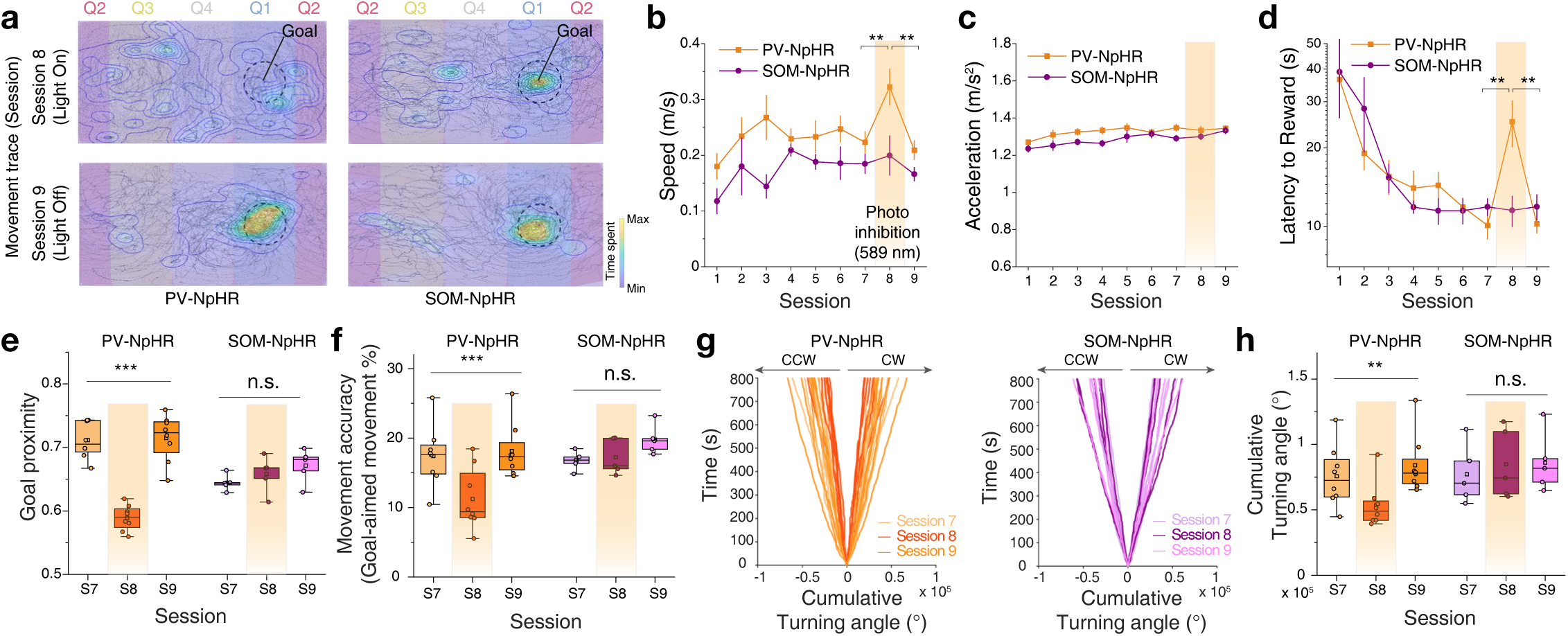
Silencing perisomatic inhibition disrupts organized motor control. **(a)** 2-dimensional projections of representative movement traces of mice on the ball in PV-NpHR and SOM-NpHR groups with photoinhibition (Session 8) and without photoinhibition (Session 9). Blockage of local glutamate release in the premotor cortex impaired organized motor control. **(b)** Average movement speed of PV-NpHR and SOM-NpHR (n = 8 mice for PV-NpHR, one-way repeated measures ANOVA with Greenhouse-Geisser correction for Sessions 7, 8, and 9, *F_session_* = 10.16, *P* = 0.0019, Fisher multiple comparisons tests, Session 7 vs. 8, *P* = 0.003; Session 7 vs. 9, *P* = 0.61, Session 8 vs. 9, *P* = 0.001; n = 5 mice for SOM-NpHR, *F_session_* = 1.08, *P* = 0.38). **(c)** Average movement acceleration (PV-NpHR, one-way repeated measures ANOVA for sessions 7, 8, and 9, *F_session_* = 0.45, *P* = 0.65; SOM-NpHR, *F_session_* = 3.69, *P* = 0.07). **(d)** Average latency to reward (PV-NpHR, one-way repeated measures ANOVA with Greenhouse-Geisser correction for sessions 7, 8, and 9, *F_session_* = 9.55, *P* = 0.014, Fisher multiple comparisons tests, Session 7 vs. 8, *P* = 0.002; Session 7 vs. 9, *P* = 0.97, Session 8 vs. 9, *P* = 0.0021; SOM-NpHR, *F_session_* = 0.094, *P* = 0.84). **(e)** Average goal proximity of PV-NpHR and SOM-NpHR in Sessions 7, 8, and 9 (PV-NpHR, one-way repeated measures ANOVA with Greenhouse-Geisser correction, *F_session_* = 40.67, *P* = 3.05 × 10^−5^, Fisher multiple comparisons tests, Session 7 vs. 8, *P* = 2.12 × 10^−6^; Session 7 vs. 9, *P* = 0.84, Session 8 vs. 9, *P* = 1.56 × 10^−6^; SOM-NpHR, *F_session_* = 1.57, *P* = 0.28). **(f)** Movement accuracy of PV-NpHR and SOM-NpHR in Sessions 7, 8, and 9 (PV-NpHR, one-way repeated measures ANOVA with Greenhouse-Geisser correction, *F_session_*= 13.9, *P* = 4.72 × 10^−4^ Fisher multiple comparisons tests, Session 7 vs. 8, *P* = 7.39 × 10^−4^; Session 7 vs. 9, *P* = 0.62, Session 8 vs. 9, *P* = 2.84 × 10^−4^; SOM-NpHR, *F_session_* = 3.06, *P* = 0.14). **(g)** Cumulative turning angle of PV-NpHR and SOM-NpHR over time. **(h)** Comparison of cumulative turning angle between PV-NpHR and SOM-NpHR (PV-NpHR, Friedman test for Sessions 7, 8, and 9, *χ*^2^*(2)* = 13, *P* = 0.0015, Dunn’s multiple comparisons tests, Session 7 vs. 8, *P* = 0.037; Session 7 vs. 9, *P* = 0.95, Session 8 vs. 9, *P* = 0.0014; SOM-NpHR, *χ*^2^*(2)* = 2.8, *P* = 0.25). n.s., not significant; \*\**P* < 0.01; \*\*\**P* < 0.001; Error bars indicate s.e.m. In the box plot, the midline, box size, and whisker indicate median, 25-75th percentile, and 10-90th percentile, respectively.

**Extended Data Figure 5.**
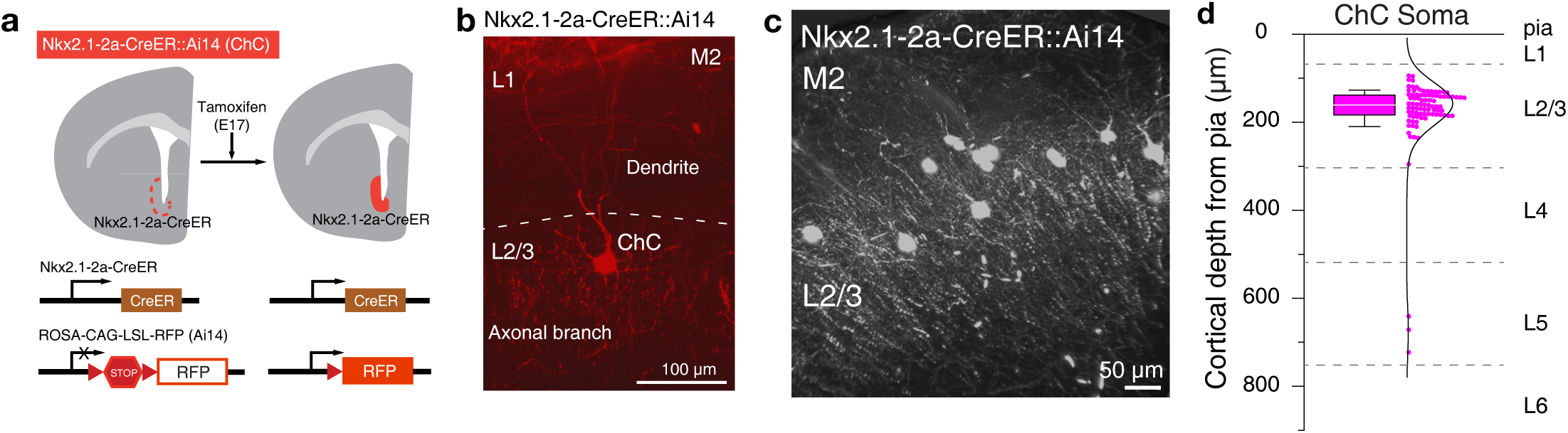
Genetical labeling of cortical chandelier cells in L2/3 premotor cortex. **(a)** Generation of Nkx2.1-2a-CreER::Ai14 mice. Nkx2.1-2a-CreER::Ai14 were generated by crossing Nkx2.1-2a-CreER with Ai14 mouse lines. A 2A-CreER cassette was inserted into the frame immediately after an open reading frame of an Nkx2.1 gene. To induce CreER activity in the offspring, tamoxifen was administered to timed pregnant SW females by oral gavage at E17. **(b)** A representative image of ChC located in layer 2/3 premotor cortex. **(c)** Light-sheet microscope image of ChCs’ densely branching axonal cartridges in layer 2/3 of the premotor cortex. **(d)** Cortical depth of ChC soma location from pia (n = 90 ChCs from 5 mice). Most of the ChCs marked by viral expression were located in the upper L2/3 (87 out of 90 ChCs) and 3 % of the ChCs were in the L5 (3 out of 90 ChCs). In the box plot, the white line, box size, and whisker indicate median, 25-75th percentile, and 10-90th percentile, respectively.

**Extended Data Figure 6.**
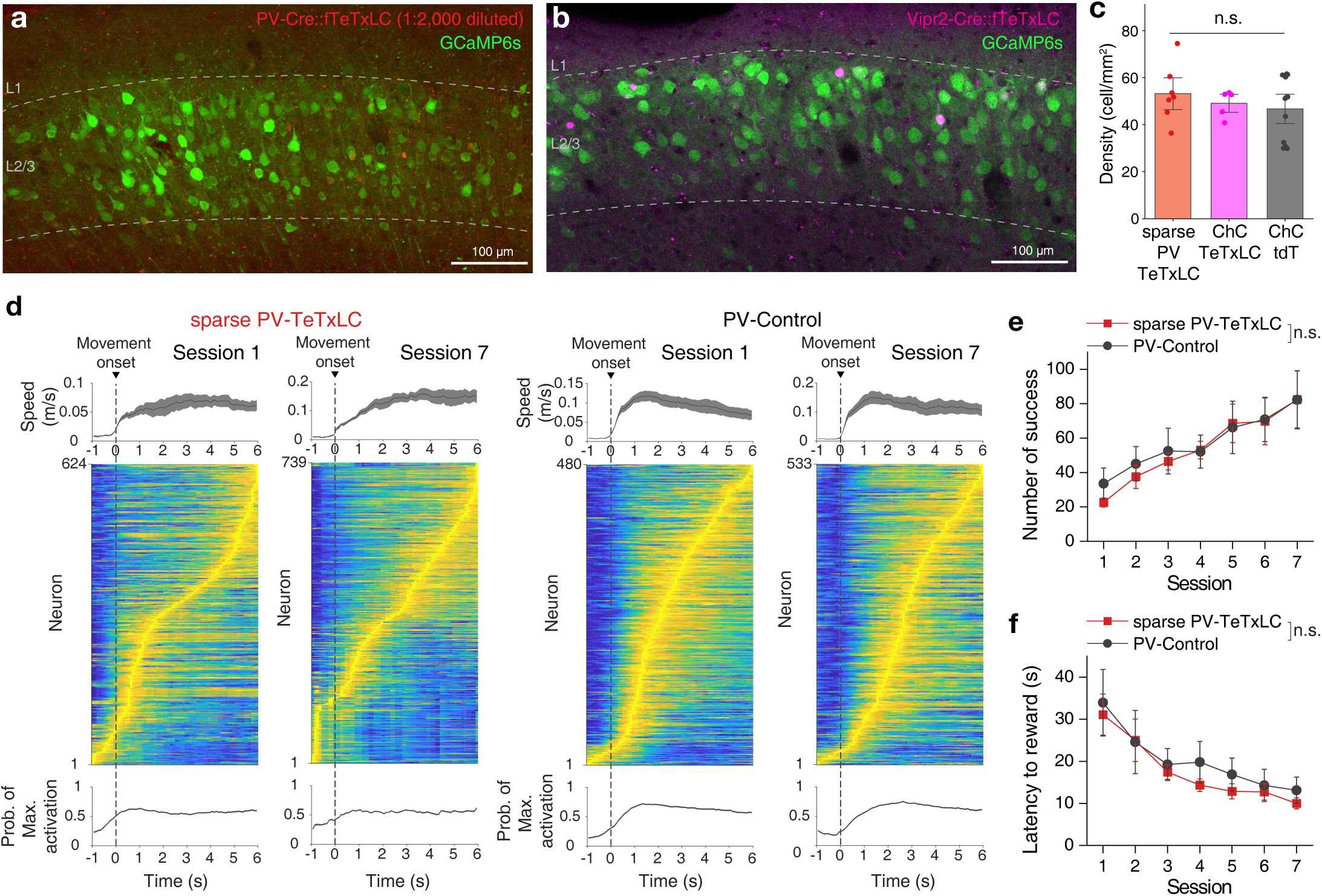
Sparse expression of tetanus toxin light chain in parvalbumin interneurons. **(a)** Example image of the expressions of AAV-hSyn-GCaMP6s (green) and AAV-hSyn-FLEX-TeTxLC-P2A-NLS-dTomato (red, 1:2,000 diluted) in L2/3 premotor cortex of PV-Cre mice (sparse PV-TeTxLC). **(b)** Example image of the expressions of AAV-hSyn-GCaMP6s (green) and AAV-hSyn-FLEX-TeTxLC-P2A-NLS-dTomato (magenta) in L2/3 premotor cortex of Vipr2-Cre mice (ChC-TeTxLC). **(c)** The density of cells expressing TeTxLC (n = 7 mice for sparse PV-TeTxLC; n= 5 mice for expression of ChC-TeTxLC; n= 11 mice for expression of FLEX-tdTomato in Vipr2-Cre mice, ChC-tdT; one-way ANOVA, *F_group_*= 0.62, *P* = 0.547). **(d)** Average movement speed (top), the normalized activity of premotor neurons (middle), and probability of maximum neuron activation (bottom) aligned to movement onset for sessions 1 and 7 in sparse PV-TeTxLC (n = 624 cells for session 1 and 739 cells for session 7, 7 mice) and PV control (n = 480 cells for session 1 and 533 cells for session 7, 7 mice). **(e)** The average number of successes for sparse PV-TeTxLC and PV control mice increased with learning (n = 7 mice for sparse PV-TeTxLC; n = 7 mice for PV control; two-way repeated measures ANOVA, *F_group_* = 0.050, *P* = 0.831). **(f)** The average latency to reward for sparse PV-TeTxLC and PV control mice deceased with learning (n = 7 mice for sparse PV-TeTxLC; n = 7 mice for PV control; two-way repeated measures ANOVA, *F_group_* = 0.346, *P* = 0.578). n.s., not significant; Error bars indicate s.e.m.

**Extended Data Figure 7.**
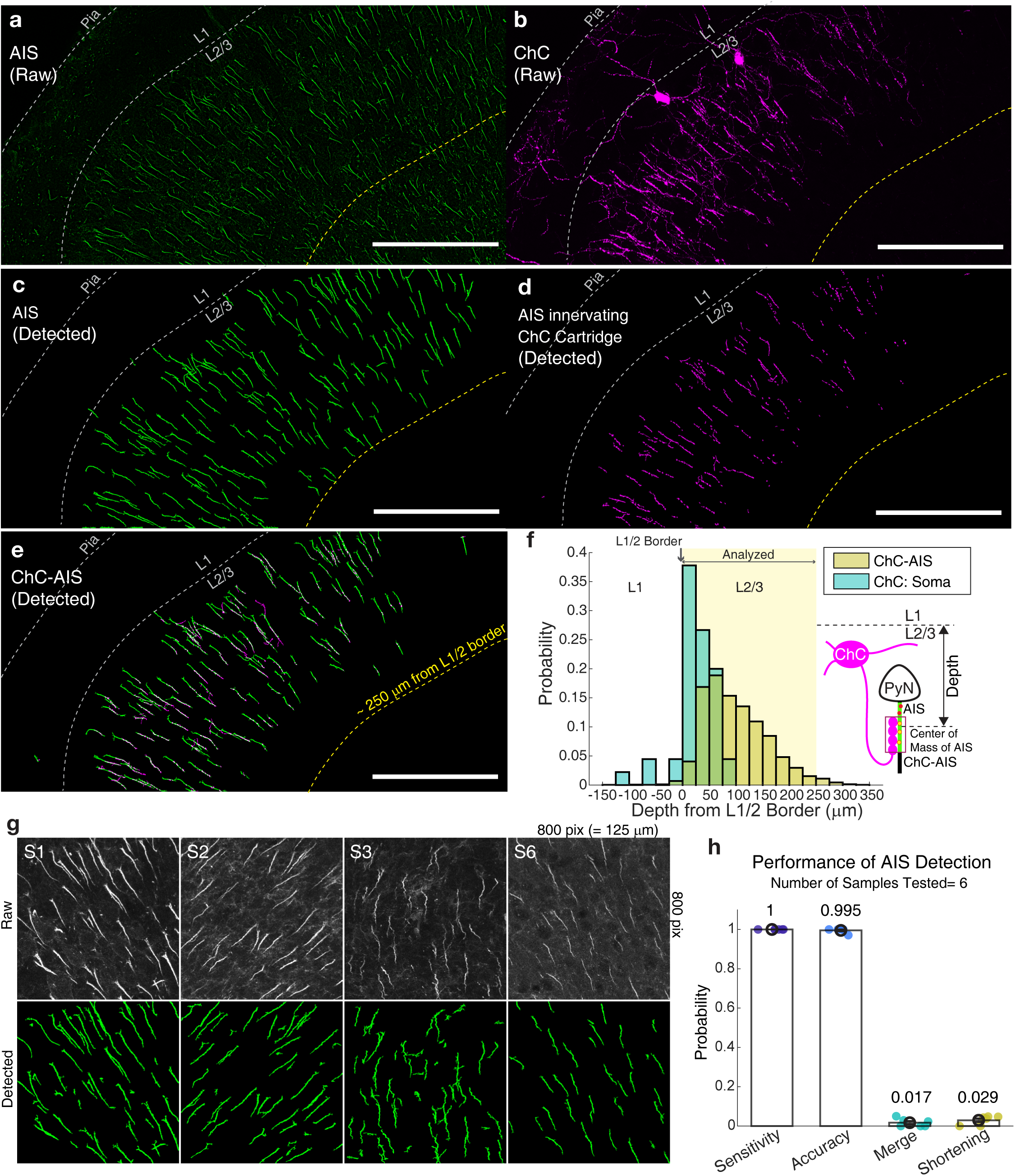
Automated detection of ChC-AIS interaction. **(a-b)** Example images of AIS and ChC in the M2 of Vipr2-Cre mice expressing AAV1-CAG-Flex-tdTomato. **(c-d)** Detected AISs and ChC axonal bouton cartridges by an automated detection tool. **(e)** Detected ChC-innervated AISs (ChC-AISs) with the white colocalized area. Representative images are maximum intensity projections of 400 X 206 X 10 μm^3^. Scale bar = 100 μm for **a-e**. The dotted lines represent the layer 1/2 border (white) and the 250 μm-deep from the border approximately (yellow). **(f)** Probability distribution of ChC somas and ChC-AISs by their position in layer 2. Only ChC-AISs within the range from 0 to 250 μm from layer 1/2 borders were considered as in layer 2 and analyzed further. **(g)** Examples of samples (125 X 125 X 10 μm^3^) used to evaluate the performance of automated AIS detection. **(h)** Performance of AIS detection evaluated by comparing the results to manual detection in 4 different categories (Sensitivity = probability that a manually detected AIS is detected automatically. Accuracy = probability that an automatically detected AIS is detected manually. Merge = probability that an automatically detected AIS corresponds to multiple AISs in manual detection. Shortening = probability that an automatically detected AIS length is significantly shorter than manual detection).

**Extended Data Figure 8.**
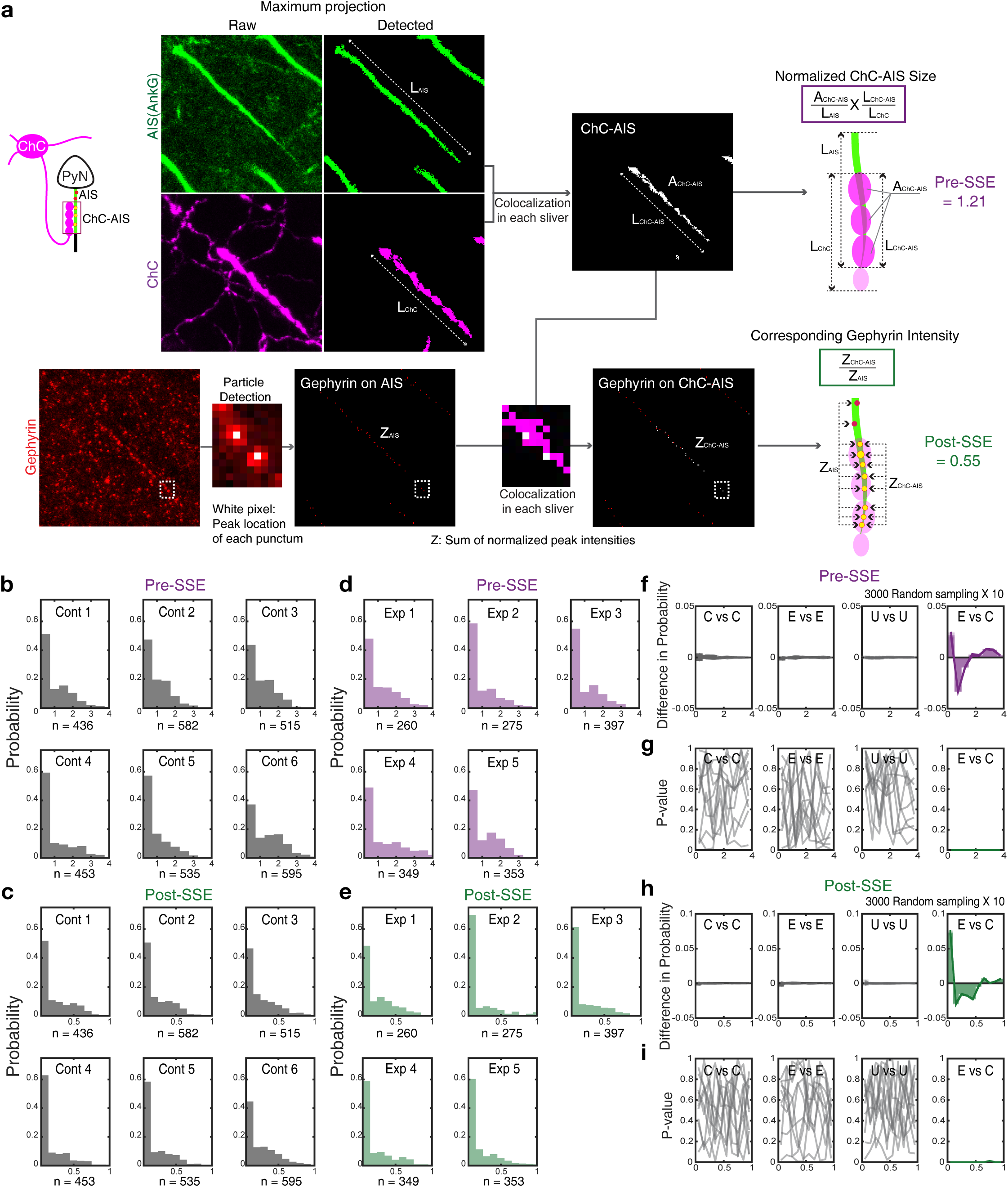
Evaluation of ChC-AIS synaptic structural efficacy. **(a)** Automatically detected segments were processed to extract quantities as the total AIS length (L_AIS_), the ChC cartridge length (L_ChC_), the AIS length covered by ChC (L_ChC_AIS_), the area of ChC on AIS (A_ChC-AIS_), and z-scored intensity of gephyrin puncta on AIS and ChC-AIS (Z_AIS_ and Z_ChC-AIS_) for characterization. Those quantities were used to compute the defined characteristic value of presynaptic structural efficacy (Pre-SSE) and postsynaptic structural efficacy (Post-SSE). Every image represents the maximum intensity projection of the corresponding volumetric stack (Details in Methods). **(b-e)** Normalized histograms of Pre-SSE and Post-SSE from mice in the control group (n = 6 mice) and the experimental (learning) group (n = 5 mice). **(f-i)** We performed a random sampling test (n = 3000) repetitively (10 times) to compare individual distributions after training. The difference in probability was significant between the Experimental and Control condition (E vs C). We did not observe any significant difference between any other pair (random sampling from the Control and Control (C vs C), Exp. vs. Exp. (E vs E), or Unlabeled vs. Unlabeled (U vs U)). The distributions were compared by a two-tailed Wilcoxon-Mann-Whitney test.

