## Extended Data Figures for "An adaptive behavioral control motif mediated by cortical axo-axonic inhibition"

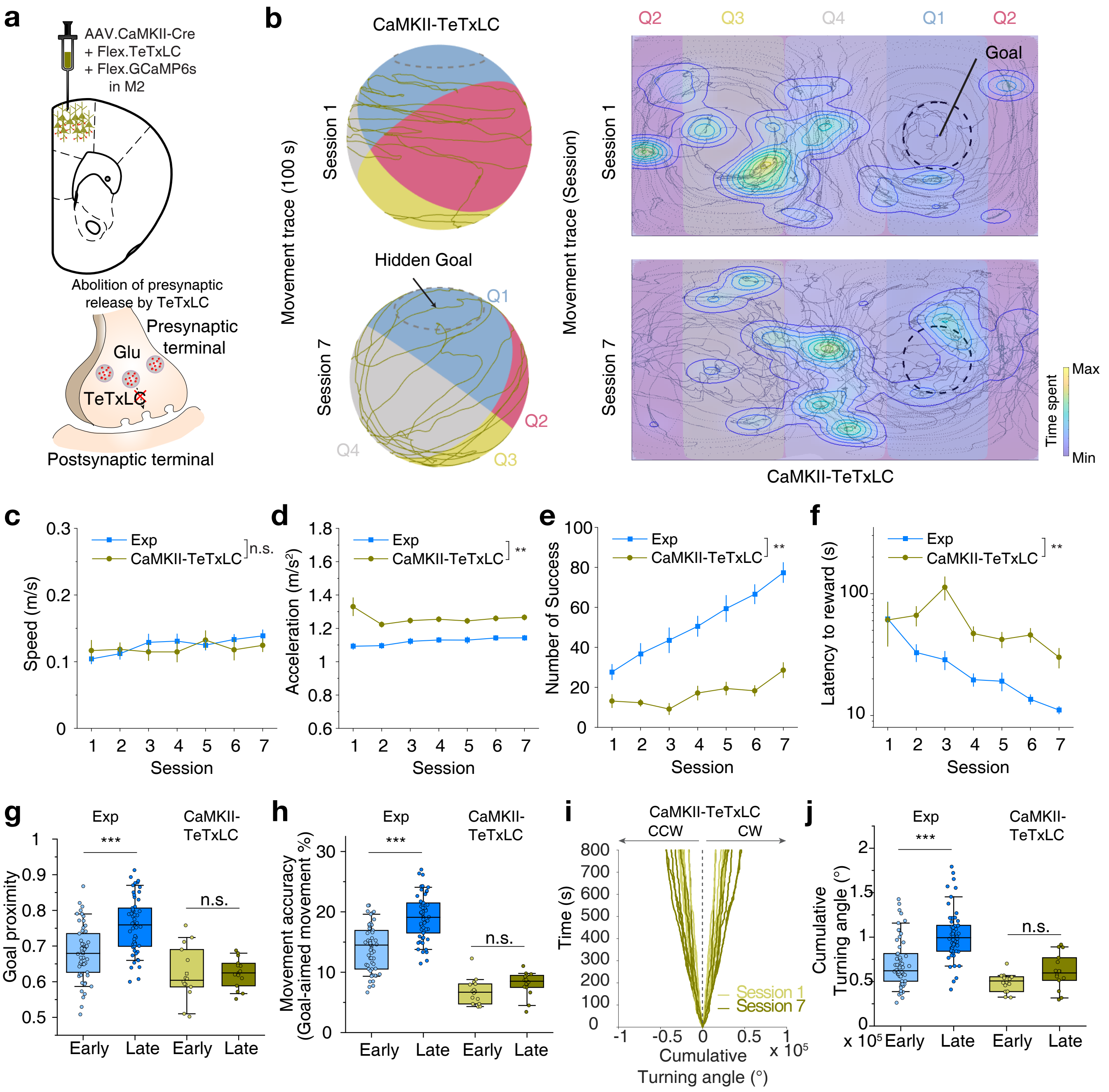


**EDF 1 - Jung et al.**

**Extended Data Figure 1. The premotor cortex is required for organized purposive motor control.**

**(a-j)** Blockage of local glutamate release in the premotor cortex impaired organized motor control.

**(a)** Schematic of virus injection of tetanus toxin light-chain, which blocks local glutamate transmission from excitatory neurons in the premotor cortex by co-expressing AAV1.CaMKII-Cre and AAV1.Flex-tetanus toxin light-chain for CaMKII-TeTxLC group.

**(b)** Representative movement trace (100 s) of a mouse in CaMKII-TeTxLC group on the ball (left) from training sessions 1 to 7 and its 2-dimensional projection (right).

**(c)** Average movement speed of CaMKII-TeTxLC (n = 27 for the experimental group, Exp; n = 7 for CaMII-TeTxLC; two-way repeated measures ANOVA, $F_{group} = 0.45$, $P = 0.53$).

**(d)** Average movement acceleration (two-way repeated measures ANOVA, $F_{group} = 19.59$, $P = 0.0044$).

**(e)** Average number of successes obtained in training sessions (two-way repeated measures ANOVA, $F_{group} = 24.11$, $P = 0.0027$).

**(f)** Average latency to reward of CaMKII-TeTxLC (two-way repeated measures ANOVA, $F_{group} = 24.11$, $P = 3.50 \times 10^{-4}$).

**(g)** Average goal proximity of CaMKII-TeTxLC (two-tailed *t*-test, $t = -5.19$, $P = 1.03 \times 10^{-6}$ for Exp; $t = -0.06$, $P = 0.95$ for CaMKII-TeTxLC).

**(h)** Movement accuracy of CaMKII-TeTxLC (two-tailed *t*-test, $t = -7.04$, $P = 2.01 \times 10^{-10}$ for Exp; $t = -1.54$, $P = 0.135$ for CaMKII-TeTxLC).

**(i)** Cumulative turning angle of CaMKII-TeTxLC over time in Sessions 1 and 7.

**(j)** Comparison of cumulative turning angle between Exp and CaMKII-TeTxLC (two-tailed *t*-test, $t = -5.84$, $P = 5.76 \times 10^{-8}$ for Exp; $t = -2.05$, $P = 0.050$ for CaMKII-TeTxLC). n.s., not significant; $^{**}P < 0.01$; $^{***}P < 0.001$; Error bars indicate s.e.m. In the box plot, the midline, box size, and whisker indicate median, 25-75th percentile, and 10-90th percentile, respectively.

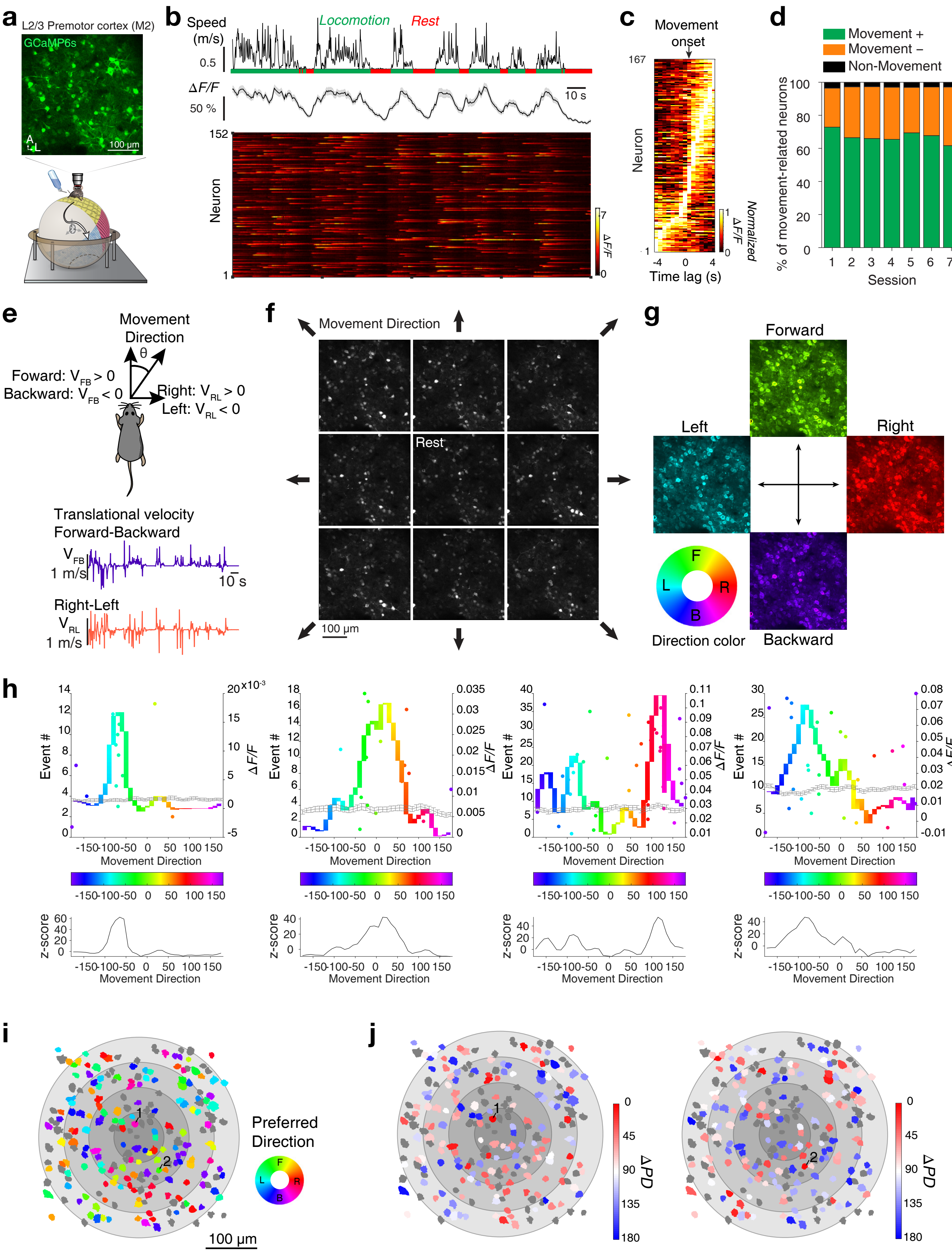


EDF 2 - Jung et al.

**Extended Data Figure 2. Movement direction selective responses of premotor neurons.**

**(a)** Schematic of the navigation task on ball maze with 2-photon calcium imaging and a representative field of view of layer 2/3 premotor neurons expressing GCaMP6.

**(b)** An example trace of mouse movement speed (top), aligned averaged fluorescence transients (middle), and a corresponding heat-map raster plot (bottom).

**(c)** Normalized fluorescence transients of premotor neurons aligned to movement onsets.

**(d)** Summary graph of movement-related neurons throughout training.

**(e)** Schematic that depicts the estimation of movement direction based on forward-backward and right-left speeds.

**(f)** Average fluorescence imaging frames during responses to varying movement directions.

**(g)** A color-coded, pixel-based map of neuron activity with respect to movement direction tuning.

**(h)** Top, four example calcium transient events by movement direction (dots) and direction tuning curves of premotor neurons (top, colored-curve). Movement direction is color-coded. The gray line indicates the average tuning curve from shuffled data. Bottom, corresponding z-scores of the actual tuning curves normalized by the tuning curves of the shuffled data.

**(i)** An exemplary preferred direction map. The color indicates the preferred direction of individual cells. Black and gray color indicate non-direction-tuned neurons. The salt-and-pepper layout of color indicates functional heterogeneity in direction-selectivity in the premotor cortex.

**(j)** Two examples of the heterogeneous spatial distribution of ΔPD (angular difference in preferred direction) between neuronal pairs. The same neuronal population as in **i**, with color-coding by ΔPD (angular difference in preferred direction between reference neuron and another direction-tuned neuron). Numbered neurons indicate reference neurons.

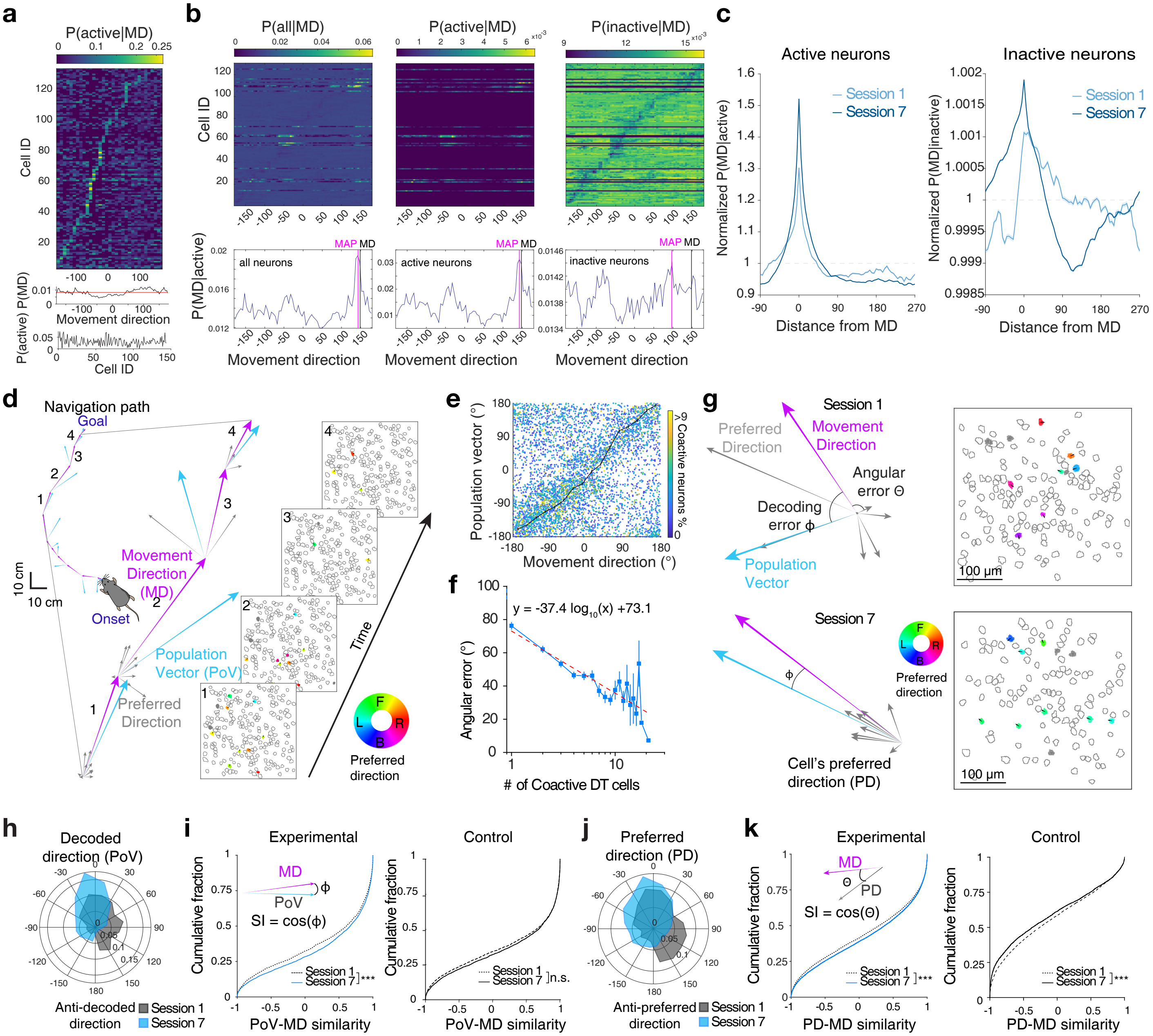


**EDF 3 - Jung et al.**

**Extended Data Figure 3. Population coding of movement direction in premotor neurons.**

**(a)** Movement direction tuning curves for each individual premotor neuron sorted from peak probability location (top). The prior probability of movement direction (middle). The marginal likelihood of being active (bottom).

**(b)** Tuning curves of neurons corresponding to movement direction at a given moment (top) and posterior probability of movement direction (bottom) given activity from all neurons (left) from active neurons (center), and from inactive neurons (left). Actual movement direction (MD) and decoded movement direction estimated with maximum a posteriori (MAP) are shown.

**(c)** An example of changes in posterior probabilities, *P(MD|A),* normalized by chance level (dotted line) from a mouse of the experiential group, for active (left) and inactive neurons (right) as a function of distance from movement direction with learning.

**(d)** Sparse population coding of movement direction during navigation; animal's movement direction (MD, magenta), preferred directions of individual active direction-tuned neurons (PD, gray), and population vector (PoV, the vector sum of the preferred directions, blue) are superimposed. Corresponding maps of active neurons during movement.

**(e)** Comparison between movement direction and sparse population vector. Each dot is color-coded by the percentage of coactive neurons in the population.

**(f)** The angular error between the population vector and actual movement direction as a function of the number of coactive direction-tuned (DT) cells. The dotted line with a negative slope coefficient indicates a linear fit of the data.

**(g)** An example of population coding of movement direction in session 1 (top) and session 7 (bottom). Magenta, cyan, and gray arrow lines indicate an animal's movement direction, population vector direction, and active neurons' preferred direction, respectively (left). Preferred direction map of the active population (right, filled ROIs). The filled color and ROI arrow indicate the corresponding preferred direction.

**(h)** Polar distribution of angular errors between PoV and MD in the experimental group (session 1, gray; session 7, light blue).

**(i)** Changes of the cumulative distribution of similarity index between PoV and MD with learning (Two-sample Kolmogorov-Smirnov test, D = 0.0495, $P$ = 0.0003 for the experimental group, n = 6 mice; D = 0.027, $P$ = 0.12 for the control group, n = 5 mice).

**(j)** Polar distribution of angular errors between PD and MD in the experimental group (session 1, gray; session 7, light blue).

**(k)** Changes of the cumulative distribution of similarity index between active neurons' PD and MD with learning (Two-sample Kolmogorov-Smirnov test, D = 0.0533, $P < 2.68 \times 10^{-10}$ for the experimental group, n = 6 mice; D = 0.0584, $P < 2.61 \times 10^{-10}$ for the control group, n = 5 mice). ***P < 0.001; n.s., not significant. Error bars indicate s.e.m.

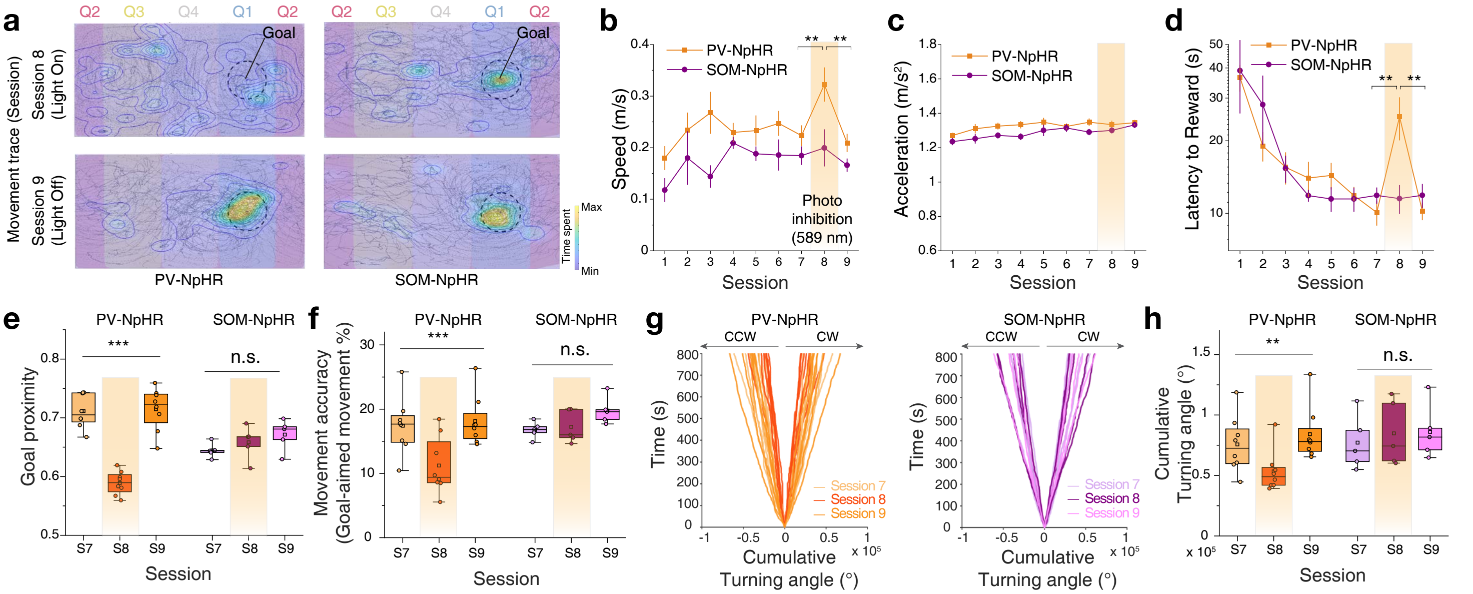


**EDF 4 - Jung et al.**

**Extended Data Figure 4. Silencing perisomatic inhibition disrupts organized motor control.**

**(a)** 2-dimensional projections of representative movement traces of mice on the ball in PV-NpHR and SOM-NpHR groups with photoinhibition (Session 8) and without photoinhibition (Session 9). Blockage of local glutamate release in the premotor cortex impaired organized motor control.

**(b)** Average movement speed of PV-NpHR and SOM-NpHR (n = 8 mice for PV-NpHR, one-way repeated measures ANOVA with Greenhouse-Geisser correction for Sessions 7, 8, and 9, $F_{session}$ = 10.16, $P$ = 0.0019, Fisher multiple comparisons tests, Session 7 vs. 8, $P$ = 0.003; Session 7 vs. 9, $P$ = 0.61, Session 8 vs. 9, $P$ = 0.001; n = 5 mice for SOM-NpHR, $F_{session}$ = 1.08, $P$ = 0.38).

**(c)** Average movement acceleration (PV-NpHR, one-way repeated measures ANOVA for sessions 7, 8, and 9, $F_{session}$ = 0.45, $P$ = 0.65; SOM-NpHR, $F_{session}$ = 3.69, $P$ = 0.07).

**(d)** Average latency to reward (PV-NpHR, one-way repeated measures ANOVA with Greenhouse-Geisser correction for sessions 7, 8, and 9, $F_{session}$ = 9.55, $P$ = 0.014, Fisher multiple comparisons tests, Session 7 vs. 8, $P$ = 0.002; Session 7 vs. 9, $P$ = 0.97, Session 8 vs. 9, $P$ = 0.0021; SOM-NpHR, $F_{session}$ = 0.094, $P$ = 0.84).

**(e)** Average goal proximity of PV-NpHR and SOM-NpHR in Sessions 7, 8, and 9 (PV-NpHR, one-way repeated measures ANOVA with Greenhouse-Geisser correction, $F_{session}$ = 40.67, $P = 3.05 \times 10^{-5}$, Fisher multiple comparisons tests, Session 7 vs. 8, $P = 2.12 \times 10^{-6}$; Session 7 vs. 9, $P$ = 0.84, Session 8 vs. 9, $P = 1.56 \times 10^{-6}$; SOM-NpHR, $F_{session}$ = 1.57, $P$ = 0.28).

**(f)** Movement accuracy of PV-NpHR and SOM-NpHR in Sessions 7, 8, and 9 (PV-NpHR, one-way repeated measures ANOVA with Greenhouse-Geisser correction, $F_{session}$ = 13.9, $P = 4.72 \times 10^{-4}$ Fisher multiple comparisons tests, Session 7 vs. 8, $P = 7.39 \times 10^{-4}$; Session 7 vs. 9, $P$ = 0.62, Session 8 vs. 9, $P = 2.84 \times 10^{-4}$; SOM-NpHR, $F_{session}$ = 3.06, $P$ = 0.14).

**(g)** Cumulative turning angle of PV-NpHR and SOM-NpHR over time.

**(h)** Comparison of cumulative turning angle between PV-NpHR and SOM-NpHR (PV-NpHR, Friedman test for Sessions 7, 8, and 9, $\chi^2(2)$ = 13, $P$ = 0.0015, Dunn's multiple comparisons tests, Session 7 vs. 8, $P$ = 0.037; Session 7 vs. 9, $P$ = 0.95, Session 8 vs. 9, $P$ = 0.0014; SOM-NpHR, $\chi^2(2)$ = 2.8, $P$ = 0.25). n.s., not significant; **$P$ < 0.01; ***$P$ < 0.001; Error bars indicate s.e.m. In the box plot, the midline, box size, and whisker indicate median, 25-75th percentile, and 10-90th percentile, respectively.

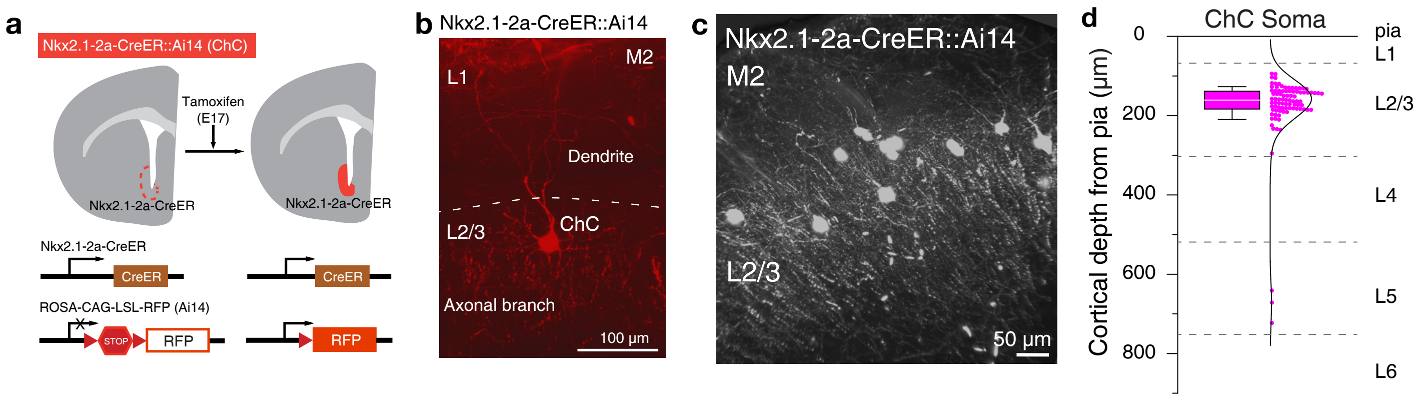


**EDF 5 - Jung et al.**

**Extended Data Figure 5. Genetical labeling of cortical chandelier cells in L2/3 premotor cortex.**

**(a)** Generation of Nkx2.1-2a-CreER::Ai14 mice. Nkx2.1-2a-CreER::Ai14 were generated by crossing Nkx2.1-2a-CreER with Ai14 mouse lines. A 2A-CreER cassette was inserted into the frame immediately after an open reading frame of an Nkx2.1 gene. To induce CreER activity in the offspring, tamoxifen was administered to timed pregnant SW females by oral gavage at E17.

**(b)** A representative image of ChC located in layer 2/3 premotor cortex.

**(c)** Light-sheet microscope image of ChCs' densely branching axonal cartridges in layer 2/3 of the premotor cortex.

**(d)** Cortical depth of ChC soma location from pia (n = 90 ChCs from 5 mice). Most of the ChCs marked by viral expression were located in the upper L2/3 (87 out of 90 ChCs) and 3 % of the ChCs were in the L5 (3 out of 90 ChCs). In the box plot, the white line, box size, and whisker indicate median, 25-75th percentile, and 10-90th percentile, respectively.

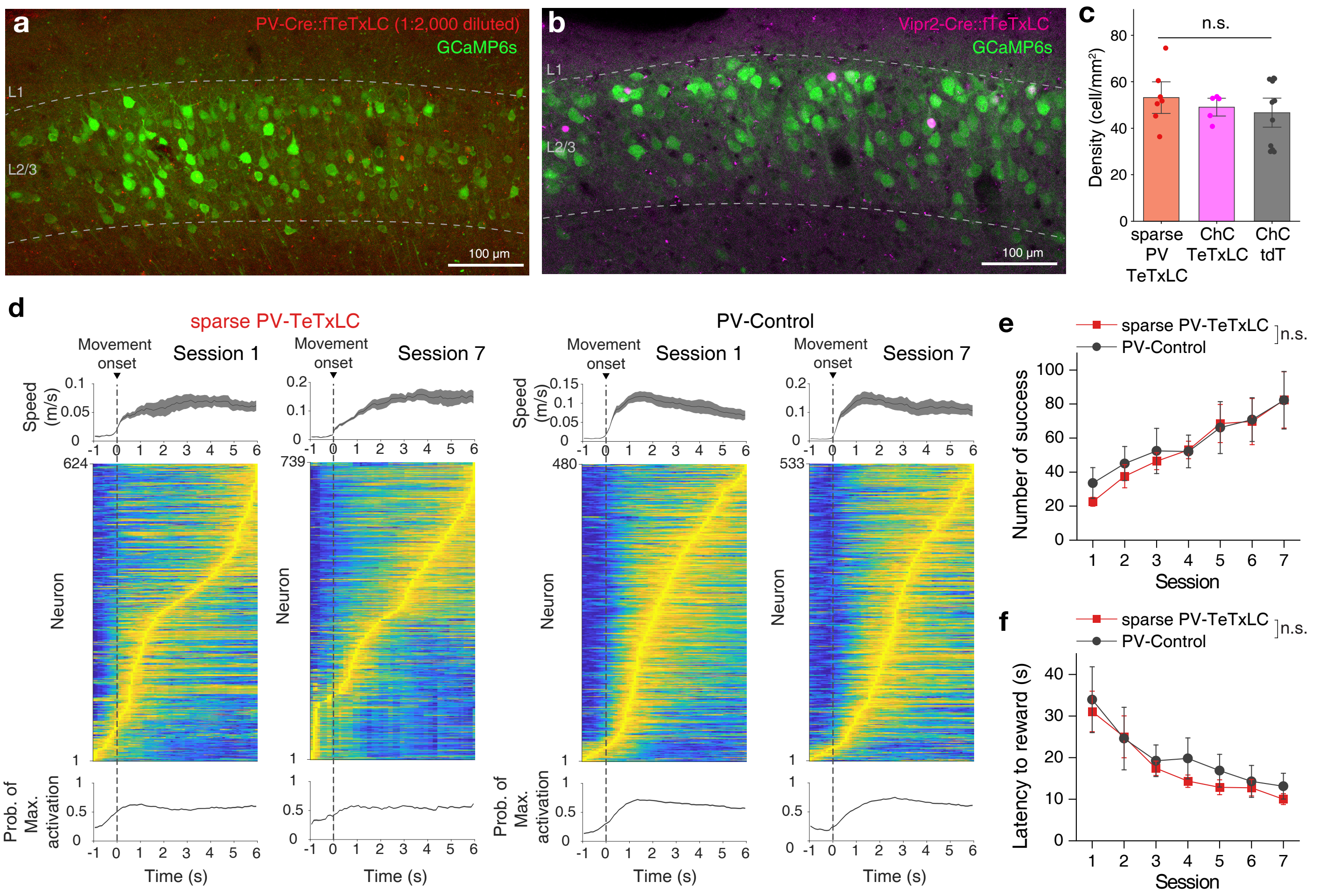


**EDF 6 - Jung et al.**

**Extended Data Figure 6. Sparse expression of tetanus toxin light chain in parvalbumin interneurons.**

**(a)** Example image of the expressions of AAV-hSyn-GCaMP6s (green) and AAV-hSyn-FLEX-TeTxLC-P2A-NLS-dTomato (red, 1:2,000 diluted) in L2/3 premotor cortex of PV-Cre mice (sparse PV-TeTxLC).

**(b)** Example image of the expressions of AAV-hSyn-GCaMP6s (green) and AAV-hSyn-FLEX-TeTxLC-P2A-NLS-dTomato (magenta) in L2/3 premotor cortex of Vipr2-Cre mice (ChC-TeTxLC).

**(c)** The density of cells expressing TeTxLC (n = 7 mice for sparse PV-TeTxLC; n= 5 mice for expression of ChC-TeTxLC; n= 11 mice for expression of FLEX-tdTomato in Vipr2-Cre mice, ChC-tdT; one-way ANOVA, $F_{group}$ = 0.62, $P$ = 0.547).

**(d)** Average movement speed (top), the normalized activity of premotor neurons (middle), and probability of maximum neuron activation (bottom) aligned to movement onset for sessions 1 and 7 in sparse PV-TeTxLC (n = 624 cells for session 1 and 739 cells for session 7, 7 mice) and PV control (n = 480 cells for session 1 and 533 cells for session 7, 7 mice).

**(e)** The average number of successes for sparse PV-TeTxLC and PV control mice increased with learning (n = 7 mice for sparse PV-TeTxLC; n = 7 mice for PV control; two-way repeated measures ANOVA, $F_{group}$ = 0.050, $P$ = 0.831).

**(f)** The average latency to reward for sparse PV-TeTxLC and PV control mice deceased with learning (n = 7 mice for sparse PV-TeTxLC; n = 7 mice for PV control; two-way repeated measures ANOVA, $F_{group}$ = 0.346, $P$ = 0.578). n.s., not significant; Error bars indicate s.e.m.

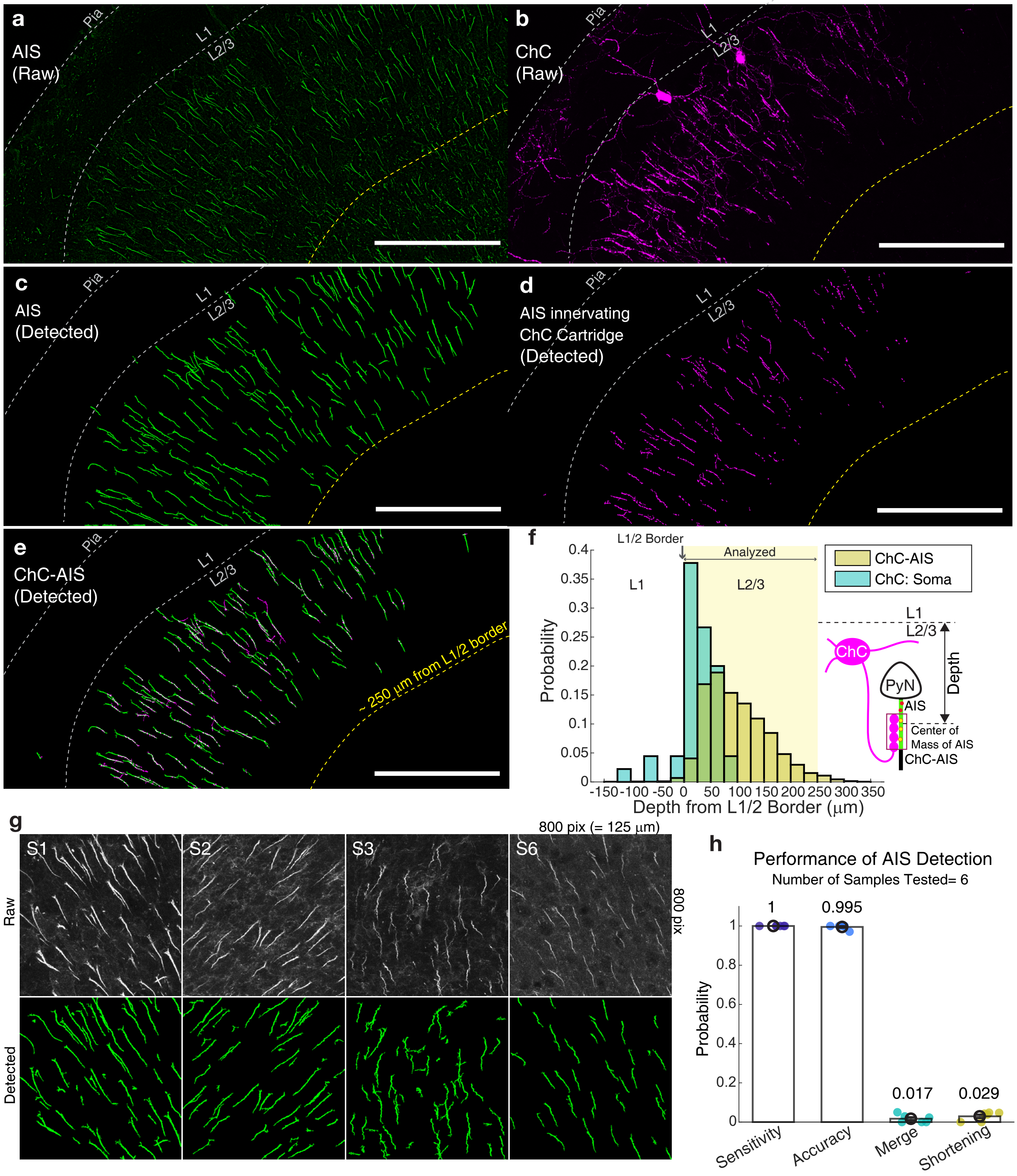


**EDF 7 - Jung et al.**

**Extended Data Figure 7. Automated detection of ChC-AIS interaction.**

**(a-b)** Example images of AIS and ChC in the M2 of Vipr2-Cre mice expressing AAV1-CAG-Flex-tdTomato.

**(c-d)** Detected AISs and ChC axonal bouton cartridges by an automated detection tool.

**(e)** Detected ChC-innervated AISs (ChC-AISs) with the white colocalized area. Representative images are maximum intensity projections of 400 X 206 X 10 $\mu m^3$. Scale bar = 100 μm for **a-e**. The dotted lines represent the layer 1/2 border (white) and the 250 μm-deep from the border approximately (yellow).

**(f)** Probability distribution of ChC somas and ChC-AISs by their position in layer 2. Only ChC-AISs within the range from 0 to 250 μm from layer 1/2 borders were considered as in layer 2 and analyzed further.

**(g)** Examples of samples (125 X 125 X 10 $\mu m^3$) used to evaluate the performance of automated AIS detection.

**(h)** Performance of AIS detection evaluated by comparing the results to manual detection in 4 different categories (Sensitivity = probability that a manually detected AIS is detected automatically. Accuracy = probability that an automatically detected AIS is detected manually. Merge = probability that an automatically detected AIS corresponds to multiple AISs in manual detection. Shortening = probability that an automatically detected AIS length is significantly shorter than manual detection).

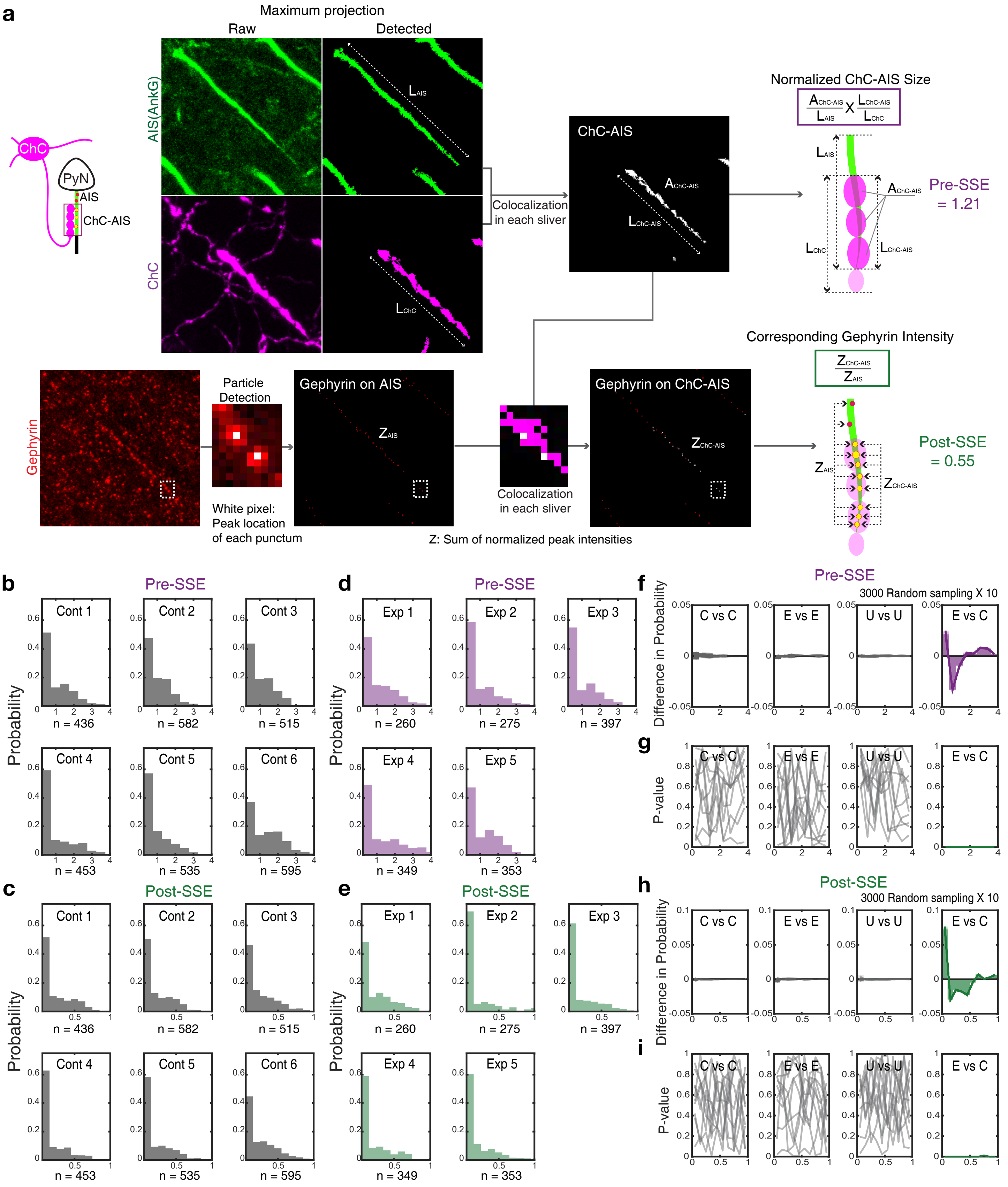


**EDF 8 - Jung et al.**

**Extended Data Figure 8. Evaluation of ChC-AIS synaptic structural efficacy.**

**(a)** Automatically detected segments were processed to extract quantities as the total AIS length ($L_{AIS}$), the ChC cartridge length ($L_{ChC}$), the AIS length covered by ChC ($L_{ChC\_AIS}$), the area of ChC on AIS ($A_{ChC\text{-}AIS}$), and z-scored intensity of gephyrin puncta on AIS and ChC-AIS ($Z_{AIS}$ and $Z_{ChC\text{-}AIS}$) for characterization. Those quantities were used to compute the defined characteristic value of presynaptic structural efficacy (Pre-SSE) and postsynaptic structural efficacy (Post-SSE). Every image represents the maximum intensity projection of the corresponding volumetric stack (Details in Methods).

**(b-e)** Normalized histograms of Pre-SSE and Post-SSE from mice in the control group (n = 6 mice) and the experimental (learning) group (n = 5 mice).

**(f-i)** We performed a random sampling test (n = 3000) repetitively (10 times) to compare individual distributions after training. The difference in probability was significant between the Experimental and Control condition (E vs C). We did not observe any significant difference between any other pair (random sampling from the Control and Control (C vs C), Exp. vs. Exp. (E vs E), or Unlabeled vs. Unlabeled (U vs U)). The distributions were compared by a two-tailed Wilcoxon-Mann-Whitney test.
